# CMAS dampens anti-tumor immunity and associates with response to neoadjuvant immunotherapy in melanoma

**DOI:** 10.64898/2026.04.27.719251

**Authors:** Magali Coccimiglio, Sofía Ibáñez-Molero, Babet O. Springer, Laura Goossens-Kruijssen, Georgia Clayton, Reece Davison, Tao Zhang, Steven Wijnen, Ernesto Rodriguez, Katarina Olesek, Sophie P. G. R. Veenstra, Thomas van den Brekel, Eleonora Nardini, Joelle van Elk, Eelco Keuning, Sara Garcia-Garcia, Kelly Boelaars, Noortje de Haan, Mariette Labots, Tanja de Gruijl, Christian Blank, Fabrizio Chiodo, Yvette van Kooyk

## Abstract

Identifying immune-regulatory pathways to predict response is crucial for the efficacy of immune checkpoint blockade (ICB) immunotherapies. Sialylation is upregulated in tumor cells and modulates immune responses in cancer, yet its impact on patient clinical outcome and the spatial organization of the tumor microenvironment remain unclear. Here, using publicly available single-cell RNA sequencing data we show that expression of the sialylation master regulator CMAS in melanoma cells correlates with poorer patient survival. Using a murine melanoma model, we demonstrate that *Cmas* deletion in tumor cells severely impaired tumor growth and improved anti-tumor lymphoid and myeloid cell responses, increasing tumor cell-intrinsic susceptibility to interferon-gamma-, CD4+ T cell-, and macrophage-mediated killing. Single-cell spatial transcriptomics on neoadjuvant ICB-treated melanoma patient tumor biopsies revealed that CMAS expression in tumor cells inversely correlated with tumor cell proximity to and activation status of T cells and macrophages. Furthermore, expression of CMAS in tumor cells was increased in patients who did not respond to immunotherapy, compared to responders. Overall, our work identifies CMAS as a key modulator of tumor-immune dynamics associated with survival and response to neoadjuvant ICB immunotherapy in melanoma patients.

## Introduction

Cutaneous melanoma was the first solid tumor in which immune checkpoint blockade (ICB) immunotherapies were tested and proven effective ^1,2^. Thus, melanoma has been a leading cancer type in ICB testing and a role model to understand the (lack of) immune responses. The treatment and outcome of melanoma patients significantly improved with neoadjuvant ICB regimes ^3,4^. However, still a fraction of patients do not show optimal responses and predicting patient outcome remains a challenge ^4,5^. Hence, a deeper understanding of mechanisms underlying immunoregulation and therapy resistance in melanoma is still needed.

Sialylation has emerged as a key regulator of immune responses in cancer ^6,7^. This pathway entails the biosynthesis of sialic acids and their addition to glycan structures present on proteins, lipids and RNA, collectively known as glycoconjugates ^6,8^. Several enzymes are responsible for the biosynthesis, activation, and transport of sialic acid to the Golgi apparatus, which constitute the donor pathway of sialylation. Subsequently, the Golgi-resident sialyltransferases catalyze the addition of the activated sialic acid donor to glycoconjugates, exposing these residues as the outermost structure on the cell surface ^6^. Modulation of tumor sialylation using metabolic inhibitors or sialidases (enzymes that cleave off surface sialic acids) increased survival and synergized with immunotherapies ^9-12^. This led to GLIMMER-01 (NCT05259696), the first phase I/II clinical trial combining ICB with an engineered human sialidase-Fc fusion in several solid tumors ^13^. However, sialic acid-containing glycoconjugates have a plethora of functions in a cell, and are involved in tumor-intrinsic cellular processes that are not yet fully understood ^14,15^.

Thus far, the sialylation pathway in human cancer has mostly been studied through single-cell RNA sequencing data ^9,11,16^. Recent advances in spatial technologies have enabled high-resolution mapping of immune cell localization within tumors, revealing strong correlations between spatial immune architectures and ICB therapy responses ^17,18^. However, how the sialylation pathway relates to the local composition and spatial organization of the tumor microenvironment (TME) *in situ* in human tissues is largely unexplored. In addition, sialylation genes have been correlated with prognosis of patients in different cancer types using bulk RNA sequencing data ^9,11,16^, but the utility of the sialylation pathway to predict patient outcome and response to ICB therapy in melanoma remains undetermined.

Here, we investigated the expression of sialylation genes in melanoma cells and found that the Cytidine Monophosphate-N-Acetylneuraminic acid Synthetase (CMAS), an enzyme that catalyzes the activation of the sialic acid donor required for all sialylation reactions, is associated with poorer survival of melanoma patients. We used a murine melanoma model to study the role of CMAS in tumor development and anti-tumor immune responses, as well as tumor-intrinsic susceptibility to immune-mediated killing. To investigate the relationship between CMAS and the spatial localization and activation of immune cells in human melanoma, we performed single-cell spatial transcriptomics and RNA sequencing on neoadjuvant ICB-treated melanoma patient samples. As we made use of samples pre- and post-ICB treatment, from responders and non-responders, we determined how *CMAS* expression correlates with ICB response.

## Materials and Methods

### Single-cell RNA sequencing analysis of benign versus melanoma tissues

The single-cell RNA sequencing dataset from Stubenvoll et al.^19^ was analysed using the package Seurat (v5.3.0, RRID:SCR_016341) in RStudio. Cells with more than 200 genes, at least 1000 counts and a maximum of 15% of mitochondrial gene expression were retained for analysis. Samples were integrated, normalized using default values and 2000 variable features were identified. Principal component analysis (PCA) was performed, followed by clustering using the first 30 PCA dimensions, resolution of 1 and Uniform Manifold Approximation and Projection (UMAP) for visualization. Clusters with less than 25 events were excluded from the analysis. For clusters in which the annotation was not clear, the function FindMarkers was used to identify them. “Low quality” clusters were defined according to the expression of markers from different cell types simultaneously. A total of 11 clusters were annotated based on known lineage markers from literature and the cell types found by Stubenvoll et al. After clustering, the expression of two sialylation gene signatures (Fig. 1A,B) was determined in melanocytes using the AddModule function.

**Figure 1.**
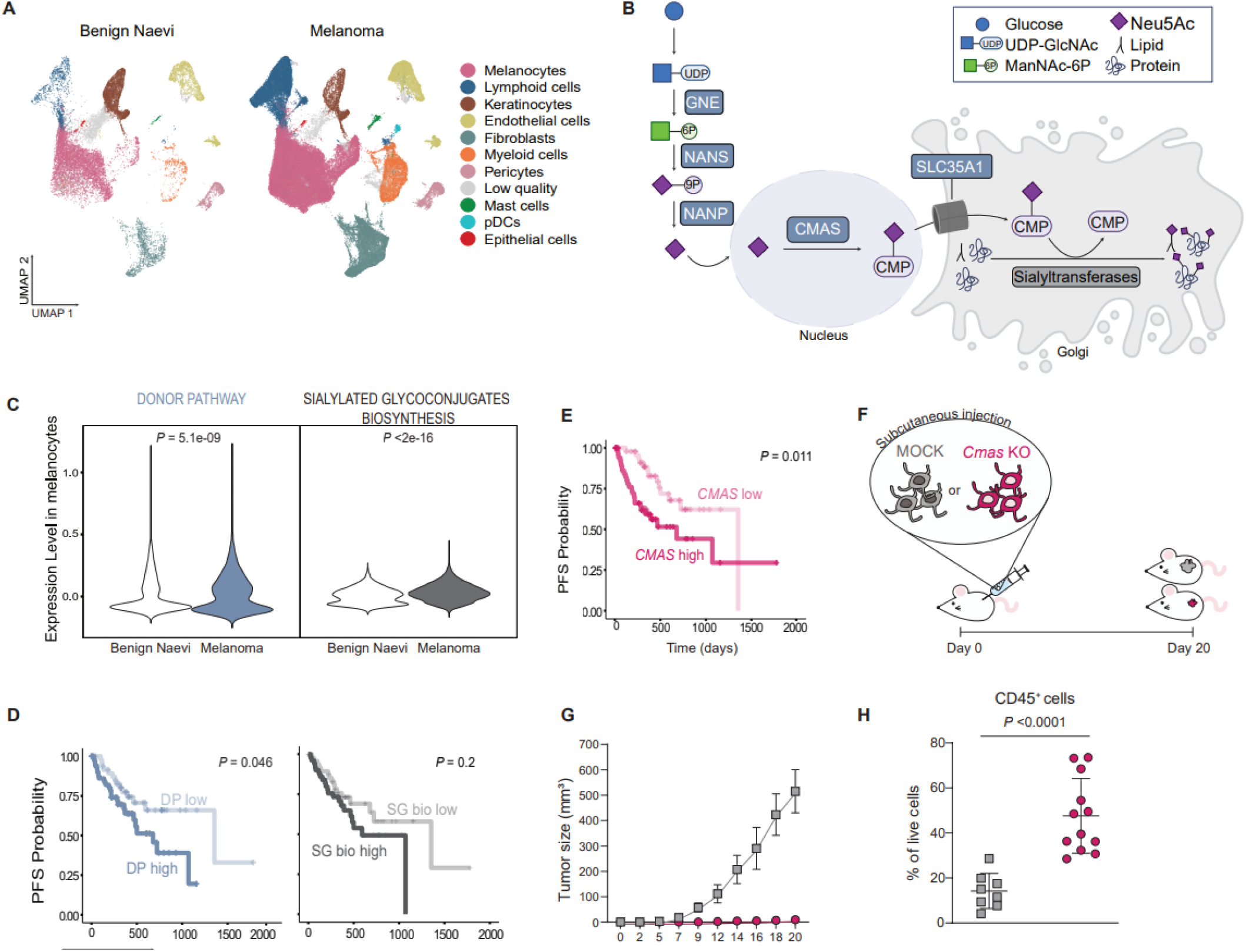
CMAS is associated with worse patient outcome and controls tumor growth and immune cell infiltration in melanoma. **A**, UMAP plots of the integrated benign naevi and primary cutaneous melanoma samples from the single-cell RNA sequencing dataset from Stubenvoll et al.^19^, coloured by cell type. **B**, Scheme of the sialylation pathway including the genes used for the donor pathway (DP) and sialylated glycoconjugates biosynthesis (SG bio) gene signatures. The DP signature included: GNE, NANS, NANP, CMAS and SLC35A1. The SG bio signature included those genes in the donor pathway signature and all sialyltransferases (ST3GAL1, ST3GAL2, ST3GAL3, ST3GAL4, ST3GAL5, ST3GAL6, ST6GAL1, ST6GAL2, ST6GALNAC1, ST6GALNAC2, ST6GALNAC3, ST6GALNAC4, ST6GALNAC5, ST6GALNAC6, ST8SIA1, ST8SIA2, ST8SIA3, ST8SIA4, ST8SIA5, ST8SIA6). Image modified from BioRender.com. **C**, Violin plots showing expression of the DP and SG bio signatures in melanocytes from benign and primary melanoma samples. Wilcoxon rank-sum test used for statistical analysis. **D**, Kaplan-Meier plots of progression-free survival (PFS) probability over time in primary cutaneous melanoma patients from TCGA with low (n=52) or high (n=51) expression of the DP and SG bio signatures, based on the median. Log-rank test used for statistical analysis. **E**, Kaplan-Meier plot of PFS over time in primary cutaneous melanoma patients from TCGA with low (n=52) or high (n=51) CMAS expression, based on the median. Log-rank test used for statistical analysis. **F**, Scheme of the in vivo tumor experiments in which B16OVA MOCK (n=18) or Cmas KO (n=38) cells were injected subcutaneously in the flank of mice. **G**, Size of B16OVA MOCK and Cmas KO subcutaneous tumors over time. Data pooled from 2 independent in vivo experiments. Data shown as mean ± s.e.m. **H**, Percentage of immune cells in B16OVA MOCK and Cmas KO tumors. Data shown as mean ± s.d. (n=8 mice in MOCK group and n=12 mice in Cmas KO group). Unpaired t-test used for statistical analysis. pDCs: plasmacytoid dendritic cells, UDP-GlcNAc: uridine diphosphate-N-acetyl glucosamine, GNE: UDP-GlcNAc 2-epimerase/ManNAc-6-kinase, ManNAc-6P: N-acetyl-mannosamine 6-phosphate, NANS: Neu5Ac 9-phosphate synthase, NANP: Neu5Ac 9-phosphate phosphatase, CMAS: CMP-Neu5Ac synthase, CMP-Neu5Ac: cytidine monophosphate N-acetyl neuraminic acid, SLC35A1: Solute Carrier Family 35 Member A1, TCGA: The Cancer Genome Atlas, s.d.: standard deviation, s.e.m.: standard error of the mean, OVA: ovalbumin

### Survival analysis of melanoma patients from TCGA

Data from The Cancer Genome Atlas (TCGA) Melanoma study was obtained using the Xena browser^20^. We used primary samples and removed samples with null data. *CMAS* gene expression, progression-free interval and time-to-event data were downloaded and Kaplan-Meier plots were made using the packages survival and survminer in RStudio.

### Generation of B16OVA MOCK and *Cmas* KO cells

The murine melanoma cell line B16OVA was obtained from Prof. T.N. Schumacher (Netherlands Cancer Institute)^21^ and cultured in RPMI-1640 medium (Gibco) supplemented with 10% Fetal Calf Serum (Biowest), 2 mM L-glutamine and 1000 U/mL Penicillin-Streptomycin (all Gibco) (RP10 medium), and incubated at 37°C and 5% CO^2^. B16OVA cells were plated in 6-wells flat bottom plates (100.000 cells/well). After 2 days, they were transfected with either MOCK CRISPR-Cas9 plasmids or the same plasmid containing guide-RNAs targeting the murine *Cmas* gene as previously described ^11,22^. Cells were sorted using the BD FACSAria™ Fusion sorter based on negative sialic acids staining (described below, Supplementary Fig. S1). B16OVA MOCK and *Cmas* KO cells were maintained in RP10 medium supplemented with 1ug/ml puromycin (Invivogen).

### Sialic acids flow cytometry staining

Cells were washed with Phosphate Saline Buffer (PBS), followed by staining with Fixable Viability Dye eFluor™ 450 (Invitrogen) for 10 minutes. After washing with Hanks’ Balanced Salt Solution + 0.5% Bovine Serum Albumin (HBSS-BSA), cells were incubated with a mix of pan-Lectenz (2ug/ml, LectenzBio) and streptavidin-APC in HBSS-BSA for 30 minutes. Cells were washed once with PBS and fixated with 2% Paraformaldehyde (PFA) in PBS for 15 minutes. Cells were washed and resuspended in PBS 0.5% BSA (PBA) for acquisition in the BD LSRFortessa X-20 cytometer. All incubations were performed at 4°C in the dark. FlowJo v10.7 was used for data analysis.

### Glycomics of B16OVA MOCK and *Cmas* KO cells

#### a) Chemicals and reagents

Ammonium hydroxide solution, glacial acetic acid, trifluoroacetic acid, formic acid (FA), 2-propanol and potassium hydroxide (KOH) were purchased from Honeywell Fluka. Supel Carbon, 2.7 μm particle PGC HPLC column (5 cm x 2.1 mm; Supel Carbon analytical column) and ethanol were purchased from Merck (Darmstadt, Germany). Ammonium bicarbonate (ABC), sodium borohydride (NaBH4), cation-exchange resin Dowex (50W-X8), DL-dithiothreitol (DTT) and hydrochloric acid (HCl) were obtained from Sigma-Aldrich (Steinheim, Germany). The GSL C17:0 isoglobotriaosylceramide (iGb3) was purchased from Avanti Polar Lipids (Brimingham, AL). Endoglycoceramidase I (EGCase I, recombinant clone derived from Rhodococcus triatomea and expressed in Escherichia coli) and 10x EGCase I buffer (500 mM HEPES, 1M NaCl, 20 mM DTT, and 0.1% Brij 35, pH 5.2) were purchased from New England BioLabs Inc. (Ipswich, MA). Peptide N-glycosidase F (PNGase F) was obtained from Roche Diagnostics (Mannheim, Germany). 8 M guanidine hydrochloride (GuHCl) was obtained from Thermo Fisher Scientific (Waltham, MA). MultiScreen HTS 96-well plates (hydrophobic Immobilon-P PVDF membrane) were obtained from Millipore (Amsterdam, the Netherlands) and 96-well PP filter plates from Orochem Technologies (Naperville, IL). Bulk sorbent Carbograph was obtained from Grace Discovery sciences (Columbia, SC). Immobilized boronic acid resin was obtained from Pierce (Product number: 20244, Thermo Fisher Scientific). Acetonitrile and methanol were purchased from Actu-All Chemicals (Oss, the Netherlands). Ultrapure water was generated from an ELGA Labwater system (Ede, the Netherlands).

#### b) Analysis of released GSL-, N-, and O-glycans using porous graphitized carbon nano-LC-ESI-MS/MS

GSL-, N-, and O-glycan alditols were prepared from 0.5 million cells in 96-well plate format as previously described with slight modification ^23,24^. In brief, cell lysates were applied to the hydrophobic Immobilon-P PVDF membrane in a 96-well plate format. Protein denaturation was achieved by applying 75 µL denaturation mix (72.5 µL 8 M GuHCl and 2.5 µL 200 mM DTT) in each well, followed by removal of the unbound material by centrifugation. GSL-glycans were firstly released using 12 mU EGCase I at 37 degrees for 16 hours. After removal of N-glycans using PNGase F, the O-glycan alditols were released from the same PVDF membrane-immobilized sample via reductive β-elimination at 50 degrees for 16 hours. GSL isoglobotriaosylceramide (iGb3, 4 ng) Maltoheptaose (DP7, 4 ng) and maltopentaose (DP5, 2 ng) were added as internal standards to support absolute quantification of GSL-glycans, N-glycans, and O-glycans, during the respective release steps. GSL-glycan and N-glycan alditols were prepared using reduction following a published procedure ^23,25^. All three types of glycans were desalted using by cation exchange desalting in a filter plate format. The desalted GSL-glycan and N-glycan alditols were further purified by PGC-SPE while desalted O-glycan alditols were purified by PGC-BOR SPE as previously described ^26^.

Glycan alditols were dissolved in 10 μL of water prior to porous graphitized carbon nano-liquid chromatography (PGC nano-LC)-ESI-MS/MS analysis. Home-packed PGC trap column (2.7 μm Hypercarb, 320 μm x 30 mm) and PGC nano-column (2.7 μm Hypercarb 100 μm x 150 mm) connected to a timsTOF felX mass spectrometer (Bruker Daltonics, Bremen, Germany) were used for the separation and detection of glycans. The temperature of the column was maintained constant at 35 degrees. Ionization was achieved using the CaptiveSpray nanoBooster source (Bruker) with isopropanol-enriched dopant nitrogen gas and a capillary voltage of 1000 V applied in negative ion mode. MS spectra were acquired within an m/z range of 300–1850 for GSL-glycans, 500–1850 for N-glycans, and 220-1850 for O-glycans, respectively. MS/MS spectra were generated using collision-induced dissociation for the top five most abundant precursors per MS spectrum covering an m/z range from 100 to 3000. Glycan structures were assigned based on the known MS/MS fragmentation patterns in negative-ion mode ^27^, elution order, and general glycobiological knowledge, with help of Glycoworkbench ^28^ and Glycomod ^29^ software. Quantification and quality control (mass accuracy, isotopic pattern matching) was performed in Skyline 21.1.0.146 (ProteoWizard) ^30^. Relative quantification of individual glycans was performed by normalizing the total peak area of all glycans within one sample to 100% and expressing each individual glycan as a percentage thereof. To estimate the glycan amount per cell, glycan intensity was normalised to the intensity of the internal standard. Afterwards, assuming the complete release of glycans and similar response factors between released glycan and standard, the number of glycans per cell was estimated.

### CellTiter-Blue® Cell Viability Assay

B16OVA MOCK and Cmas KO cells were plated in triplicates in a 96-wells flat bottom plate. After 48 hours, the CellTiter-Blue® Cell Viability Assay (Promega) was performed according to manufacturer’s instructions. Fluorescence (ex/em=560/590nm) was measured in the BioTek Synergy HTX Multimode Reader (Agilent).

### *In vivo* experiments

C57Bl6/J mice were purchased from Charles River and kept at the Animal Research Institute Amsterdam (ARIA) of the Amsterdam UMC (The Netherlands) under pathogen-free conditions. Mice were acclimatized for 2 weeks before the start of experiments at the age of 6-8 weeks. All experiments were approved by the Animal Welfare Body from the Amsterdam UMC (IvD, AVD11400202317179, SP2400567). Mice were injected subcutaneously in the flank with B16OVA MOCK or *Cmas* KO cells (300.000 cells/100ul PBS) under 2-3% isoflurane. Tumor growth and animal welfare were monitored 3 times per week. A calliper was used to measure length (L) and width (W) of the tumor, and size was determined with the formula: π*(L*W*((L+W)/2))/6.

### Sialic acids tissue staining (immunohistochemistry)

Tumor tissues were harvested in Cryomatrix™ embedding resin (Epredia) and snap frozen in liquid nitrogen for subsequent cryosection at 5µm thickness. Tissue slides were stored at −20 degrees. For the staining, slides were rehydrated and blocked with Carbo-free blocking solution (Vector Labs) for 30 minutes at room temperature, followed by a 2 hours incubation with a mix of pan-Lectenz (Lectenz Bio, 4ug/ml) and streptavidin-horseradish peroxidase (HRP). Then, slides were washed with PBS and developed with DAB (DAKO). Slides were stained with haematoxylin and dehydrated, and finally mounted with Entallan for brightfield imaging on Vectra Polaris (PerkinElmer).

### Tumor immune profiling

#### a) Tumor tissue digestion

Tumors were harvested in cold RP10 medium and transferred to wells containing Liberase TL (Roche) diluted in PBS (0.75mg/ml). After cutting them into small pieces with sterile scissors, they were transferred to tubes containing the Liberase solution, and incubated 25 minutes at 37°C and shaking. Then, RP10 HE medium (RP10 medium + 10mM EDTA + 20mM HEPES + 20uM 2-mercaptoethanol) was added and they were incubated for 10 minutes at 4°C and shaking. Finally, the digested tissues were filtered through 100µm filters, spun down and resuspended in RP10 medium for subsequent counting of live cells using trypan blue and flow cytometry staining as explained below.

#### b) Profiling by spectral flow cytometry

Two different antibody panels were used to profile B16OVA *Cmas* KO and MOCK tumors (Supplementary Table S1, Supplementary Fig. S2,3). For panel 1, cells were stained with LIVE/DEAD™ Fixable Blue (Invitrogen) in PBS for 15 minutes, followed by 30 minutes incubation with a mix of antibodies to stain cell surface molecules together with True-Stain Monocyte Blocker™ and Purified anti-mouse CD16/32 Antibody (Fc block) (both Biolegend). Then, a fixation and permeabilization step was done using the Foxp3/Transcription Factor Staining Buffer Set (Invitrogen), to finally stain intracellular markers FoxP3 and Ki67 with a 30 minutes incubation. For panel 2, cells were stained with LIVE/DEAD™ Fixable Blue (Invitrogen) in PBS for 15 minutes and then 30 minutes with a mix of pan-Lectenz (2ug/ml, LectenzBio) and streptavidin-AF647, followed by 30 minutes incubation with a mix of antibodies to stain surface molecules and fixation with 4% PFA. Incubations were performed at 4°C in the dark unless stated otherwise. SpectroFlo® software was used for the unmixing of spectral flow cytometry data, and then it was uploaded into the OMIQ software for scaling and removing of low quality events using PeacoQC ^31^.

For B16OVA WT tumors, cells were stained with LIVE/DEAD™ Fixable Blue (Invitrogen) in PBS for 15 minutes. After washing with Hanks’ Balanced Salt Solution + 0.5% Bovine Serum Albumin (HBSS-BSA), cells were incubated with a mix of pan-Lectenz (2ug/ml, LectenzBio) and streptavidin-BUV615 (Becton Dickinson) in HBSS-BSA for 30 minutes. This was followed by a wash with PBA and 30 minutes incubation with a mix of antibodies to stain cell surface molecules (Supplementary Table S2) together with True-Stain Monocyte Blocker™ and Purified anti-mouse CD16/32 Antibody (Fc block) (both Biolegend).

SpectroFlo® software was used for the unmixing of spectral flow cytometry data. FlowJo v10.7 was used for further data analysis.

### Tumor killing and MHC expression upon IFN-γ stimulation

B16OVA MOCK and *Cmas* KO cells were seeded in a 96-well flat bottom plate (25.000 or 3.000 cells/well, for overnight or 3-days incubation, respectively) in RP10 medium alone or containing variable concentrations of mouse IFN-γ (ImmunoTools), and incubated at 37°C and 5% CO^2^.

For MHC-II expression, after overnight incubation cells were washed with PBS and stained with Fixable Viability Dye eFluor™ 450 (Invitrogen) in PBS for 15 minutes. After washing the cells with PBA, they were incubated 30 minutes with anti-mouse MHC Class II (I-A/I-E) Monoclonal Antibody (M5/114.15.2) APC-eFluor™ 780 and MHC Class I (H-2Kb) Monoclonal Antibody (AF6-88.5.5.3) PE (both Invitrogen). Cells were washed with PBS and fixed with 2% paraformaldehyde (PFA) in PBS for 15 minutes. Cells were resuspended in PBA for acquisition in the BD LSRFortessa X-20 cytometer. All incubations were performed at 4°C in the dark. FlowJo v10.7 was used for data analysis.

For IFN-γ-mediated killing, the CellTiter-Blue® Cell Viability Assay (Promega) was performed according to manufacturer’s instructions. Tumor cells alone were considered as 100% viability, and the percentage viability with each condition was calculated. The % killing was determined as 100 - %viability for each condition.

### T cell-mediated killing assay

#### a) Isolation of OT-I and OT-II T cells

OT-I and OT-II C57Bl6/J mice were bred in the ARIA of the Amsterdam UMC (The Netherlands) under pathogen-free conditions. Spleens from these mice were mashed through a 70µm filter in a tube containing T cell medium (RP10 medium + 50uM 2-mercaptoethanol). After spinning down, the pellet of splenocytes were resuspended in T cell medium. OT-I and OT-II splenocytes were primed 3 days in the presence of in-house produced OVA257-264 OT-I peptide (SIINFEKL, 1ug/ml) or OVA323-339 OT-II peptide (ISQAVHAAHAEINEAGR, 100ug/ml), respectively. At day 3, primed OT-I CD8 and OT-II CD4 T cells were isolated using the MagniSort™ Mouse T cell Enrichment Kit (Thermo Fisher) according to the manufacturer’s instructions. T cells were kept in T cell medium supplemented with mouse IL-2 (ImmunoTools, 50U/ml) for 2-4 hours until further use

#### b) Tumor-T cell co-cultures

B16OVA MOCK and *Cmas* KO cells were harvested and plated in a 96-wells plate in triplicates (25.000 cells/well). Variable numbers of primed OT-I CD8 or OT-II CD4 T cells were added to the tumor cells. After incubation overnight, CellTiter-Blue® Cell Viability Assay was performed and % killing was calculated as described before.

### Tumor-Bone Marrow-Derived Macrophages co-cultures

#### a) BMDM generation

Bone marrow derived macrophages (BMDM) were obtained from C57Bl6/J mice. In short, femur and tibias were harvested and cleaned removing all tissues from them and dipping them in 70% ethanol and PBS. The end of the bones were removed by sterile scissors and using a 10ml syringe with a 25G needle RP10 medium was injected to wash the bone marrow over a 70µm filter in a tube containing RP10 medium. After centrifugation, the pellet was resuspended in RP10 medium supplemented with 15% L929 Conditioned Medium (LCM, produced in-house, containing M-CSF) and plated in petri dishes. After 3-4 days incubation at 37 degrees and 5% CO2, more RP10 medium + 15% LCM was added. At day 7, BMDM were harvested and polarized for 24 hours in RP10 medium + 5% LCM with mouse IFN-γ (ImmunoTools, 100U/ml) and LPS (100ng/ml) (pro-inflammatory macrophages) or with mouse IL-4 (ImmunoTools, 20ng/ml) (anti-inflammatory macrophages)

#### b) Phagocytosis assay

B16OVA MOCK and *Cmas* KO cells were washed and resuspended in PBS, and stained with CellTrace™ Violet Cell Proliferation Kit for flow cytometry (Invitrogen) according to manufacturer’s instructions. 100.000 labelled tumor cells were added to 100.000 BMDM in a 24-wells flat bottom plate and incubated for 4 hours. All cells were harvested and washed with PBS for staining with Fixable Viability Dye eFluor™ 780 (Invitrogen) for 15 minutes. After a wash with PBA, cells were stained with a mix of Alexa Fluor® 700 anti-mouse CD45 Antibody, True-Stain Monocyte Blocker™ and Purified anti-mouse CD16/32 Antibody (Fc block) (all Biolegend) in PBA for 30 minutes. Cells were washed with PBS and fixed with 2% PFA in PBS for 15 minutes. Cells were resuspended in PBA for acquisition in the BD LSRFortessa X-20 cytometer. All incubations were performed at 4°C in the dark. FlowJo v10.7 was used for data analysis.

#### c) BMDM activation assay

B16OVA MOCK and *Cmas* KO cells were irradiated (50 Gy) and added on top of BMDM (100.000 cells in a 24-wells flat bottom plate) and incubated for 3 days. All cells were washed with PBS and stained with Fixable Viability Dye eFluor™ 780 (Invitrogen) in PBS for 15 minutes. After a wash with PBA, cells were stained with a mix of antibodies (Supplementary Table S3) with True-Stain Monocyte Blocker™ and Purified anti-mouse CD16/32 Antibody (Fc block) (both Biolegend) in PBA for 30 minutes. Cells were washed with PBS and fixed with 2% PFA in PBS for 15 minutes. Finally, cells were resuspended in PBA for acquisition in the 4 Lasers Cytek Aurora™ cytometer. All incubations were performed at 4°C in the dark. SpectroFlo® software was used for the unmixing of spectral flow cytometry data. FlowJo v10.7 was used for further data analysis.

### Single-cell spatial transcriptomics

#### a) Patient tissue collection

Formalin-Fixed Paraffin Embedded (FFPE) blocks of tissues from patients diagnosed with resectable, macroscopic stage III melanoma who received neoadjuvant ICB with available pre- and on-treatment tissues were obtained from Amsterdam UMC, department of Medical Oncology. Patients of interest were retrospectively identified from the electronical medical records from Amsterdam UMC, department of Medical Oncology. For this research archival tissue was used, collected within the context of routine clinical practice procedures on which the Dutch Medical Research Involving Human Subjects Act does not apply. All patients treated at Amsterdam UMC have the possibility to opt-out for the use of their data and tissue for research purposes. Upon completion of lymph node dissection, pathological response had been assessed locally according to International Neoadjuvant Melanoma Consortium (INMC) guidelines. Two patients with major pathological response (MPR) (1 complete and 1 near-complete pathologic response, treated with ipilimumab plus nivolumab or pembrolizumab, respectively), herein classified as responders, and two non-responding patients (1 with pathologic non-response to ipilimumab plus nivolumab and 1 with pathologic non-response to pembrolizumab) were selected for further analysis.

#### b) Processing of tissues

Biopsies were sectioned to 4 μm thickness. RNA was extracted as per the manufacturer’s instructions (RNeasy FFPE kit, Qiagen) and quality was assessed on the Tapestation (4200; Agilent) using the High Sensitivity RNA ScreenTape kit (Agilent). Sections were further cut to remove necrotic tissue previously identified with H&E staining by a pathologist, and regions of interest were placed into a 37 degrees water bath then onto the Xenium gene expression slides (10x Genomics). Sections were stained with pre-designed probes (Human Immuno-oncology panel, 380 Genes) and custom designed probes (1-50 Add-on panel) manufactured by 10x Genomics. Staining was performed according to manufacturer’s guidelines (CG000749 | Rev B, 10x Genomics). Sections were deparaffinized and rehydrated in xylene-ethanol series and chemically de-crosslinked (CG000578 | Rev E, 10x Genomics). Padlock probes were added to the tissue and incubated overnight on the thermocycler for hybridization to RNA targets. Probes were ligated and amplified via continuous rolling cycle amplification. Additional 10x cell segmentation module was added, morphology markers targeting the cell membrane, 18s ribosomal RNA and cytoskeletal proteins were incubated overnight (CG000750 | Rev B, 10x Genomics). Tissue autofluorescence was chemically quenched and nuclear staining with DAPI carried out. After staining, samples were loaded on Xenium analyser, 15 cycles of detection probe addition, imaging and fluorescence quenching take place (CG000584 | Rev G, 10x Genomics). After imaging, data was processed by Xenium onboard analysis, probe signals were decoded and pre-processed for later downstream analysis.

#### c) Analysis of single-cell spatial transcriptomics

Cell transcriptomics and segmentation was performed with standard Xenium pipeline. Data was loaded in R using Seurat v5.3 (RRID:SCR_016341). Cells expressing less than 100 nCounts and less than 50 nFeatures were filtered. All individual Seurat objects from the different patient slides were integrated and followed standard normalization, PCA, UMAP and cluster identification analysis. We manually annotated the clusters by using lineage markers included in the Immuno-oncology panel of Xenium. Clusters annotated as the same cell type were merged. We apply re-clustering for those clusters that were shown to be spatially mixed in the UMAP (mainly clusters: “T cell mixed”, “Fibroblast”, “Tumor”, “Dendritic cell” and “Macrophages”). If a subcluster had characteristic of another cell type, we re-annotated it. The cluster “Low quality” and “Mixed”, containing low expression and mixed expression of lineage markers, respectively, were removed from further analysis. The analysis of the integrated and annotated Seurat object was performed using Seurat v5.3.0, SeuratObject v5.2.0, pheatmap package v.1.0.13, ggplot2 v3.5.2, glmGamPoi v1.18.0, arrow v21.0.0.1

The correlation of CMAS expression with the distance to a given cell type was calculated using our function NN_Single. This function calculates per individual object, per individual “Expression cell” the distance to the closest “Target cell”, as well as the expression of gene of interest in the “Expression cell”. It obtains the tissue coordinate of “Expression cell” and “Target cell” and calculates the distance using get.knnx function from FNN v1.1.4.1. Results are plotted using ggplot2 v3.5.2.

### Chromium next GEM single-cell fixed RNA profiling of formaldehyde fixed & paraffin embedded (FFPE) Tissue Sections

#### a) Patient tissue collection

Single-nucleus sequencing was performed on tumor samples from ICB treated patients from the OpACIN-neo trial (Arm B; NCT02977052, n=3) and its extension cohort PRADO (NCT02977052, n=1). These studies evaluated the feasibility of neoadjuvant combination immunotherapy (ipilimumab 1mg/kg and nivolumab 3mg/kg) in patients with resectable stage III melanoma. Patients with ≤10% viable tumor cells in the resection specimen were classified as having a MPR, which serves as a surrogate marker for event-free survival (EFS). Archived tissue microarrays (TMAs) from FFPE baseline lymph node biopsies were used to generate single-nucleus RNA-seq datasets. Two TMA blocks, each containing biopsies from nine patients, were processed.

#### b) Sample preparation: nuclei isolation from formaldehyde fixed & paraffin embedded (FFPE)

Due to the limited tissue available from these small and previously used cores, sufficient material for 10x Genomics CellPlex multiplexing was obtained for four patients. Out of 9 FFPE tissue cores present in a Tissue Microarray (TMA), 4 tissue cores were selected for single-cell fixed RNA profiling. For each sample, 7-12 sections (25 um in thickness) of 1.5 mm in diameter tissue core were collected in a 1.5 ml DNA LoBind Eppendorf tubes in order to isolate sufficient amount of nuclei for the single nuclei fixed RNA sequencing. Nuclei isolation was performed, starting with FFPE tissue deparaffinization followed by a pestle based FFPE tissue dissociation using a Liberase TH enzyme mix and sample filtration trough 30 um Miltenyi pre-separation filters, all according to protocol CG000632 of 10x Genomics. The final nuclei suspensions were counted using Propidium Iodide staining solution (Invitrogen, REF# 3566) and the LUNA-FX7™ Automated Cell Counter (Logos Biosystems).

#### c) Sample hybridization

Single nuclei gene expression analysis was performed using the Chromium Next GEM single Cell Fixed RNA profiling assay (10x Genomics, PN-1000475), according to manufactures protocol CG000527. For each sample up to 200.000 nuclei were used as input for the barcoded probe hybridization using the Human Transcriptome Probe kit (10x Genomics, PN-1000420). After probe hybridization the barcoded samples were pooled and washed according to the “Pooled Wash Workflow” option in manual CG000527. Finally, the nuclei were counted using Propidium Iodide staining solution (REF# 3566, Invitrogen) and the LUNA-FX7™ Automated Cell Counter (Logos Biosystems).

#### d) Chromium next GEM single-cell fixed RNA gem generation

For GEM generation the nuclei sample pool was loaded over 2 wells of Next GEM microfluidic Chip Q to target 60.000 nuclei in total for all 4 samples. After loading Chip Q and running it on the Chromium X, GEMs were recovered and processed to construct indexed sequencing libraries. Both libraries were quantified on a 2100 Bioanalyzer Instrument following the Agilent Technologies Protocol (Agilent DNA 7500 kit, G2938-90024).

#### e) Sequencing

A Sequence library pool was composed and quantified by qPCR, according to the KAPA Library Quantification Kit Illumina® Platforms protocol (KR0405, KAPA Biosystems). Illumina Novaseq 6000 paired-end sequencing was performed using 28 cycles for Read 1, 10 cycles for each Read i7 and i5 and 90 cycles for Read 2, using the NovaSeq 6000 SP Reagent Kit v1.5 (100 cycles) (20028401, Illumina).

#### f) Analysis of chromium next GEM single-cell fixed RNA profiling

The CellPlex experiment was pre-processed with Cell Ranger Multi (v8.0.1) using the Chromium Human Transcriptome Probe Set v1.0.1 (GRCh38-2020-A). Filtered feature–barcode matrices were subsequently analyzed in R (v4.4.3) using Seurat (v5.3.0). Initial quality control and filtering were performed on each patient sample separately prior to dataset integration. Cells were filtered based on the total number of UMIs, number of detected features, and mitochondrial gene fraction. Normalization was performed with SCTransform, followed by dimensionality reduction using PCA. The first 30 principal components were used to compute a UMAP embedding. Cell type assignment employed a consensus strategy combining automated annotation with SingleR (v2.6.0) and analyses of cluster-specific marker genes. When both methods converged, the resulting annotation was assigned. In cases where SingleR and marker-based annotation differed in the level of granularity (e.g., distinguishing monocyte-derived dendritic cells from plasmacytoid dendritic cells), we prioritized broader classifications (e.g., “dendritic cell”) unless unambiguous evidence supported a more specific subtype. If no clear consensus could be reached, manual curation was performed by considering marker expression patterns and, if present, by looking at closely related clusters in the UMAP embedding. Ultimately, only two clusters from a single patient remained unclassified with certainty and were labeled as unidentified Melanoma and Unidentified Lymphocyte. After per patient pre-processing, we merged the datasets and ended up with a total of 66.203 cells. DGE analyses between the Malignant cells of the two MPR and two non-MPR patients was performed.

### Statistical analysis

Statistical analysis was performed using GraphPad Prism 10 or RStudio. The statistical tests used are mentioned in the figure legends. P values are depicted in the figures and statistical significance was considered when P<0.05.

### Data availability

The single-cell RNA sequencing dataset from Stubenvoll et al.^19^ was obtained from NCBI GEO database (accession number: GS277165). The data generated in this study are available upon request from the corresponding author. Code for the analysis of single-cell spatial transcriptomics and Chromium Next GEM Single-cell fixed RNA profiling will be made available at http://github.com.

## Results

### CMAS is associated with worse patient outcome and controls tumor growth and immune cell infiltration in melanoma

To investigate whether changes in sialylation gene signatures are related to the malignant transformation of melanocytes, we analyzed single-cell RNA sequencing data that include both benign naevi and primary cutaneous melanoma tissues ^19^ (Fig. 1A, Supplementary Fig. S4A-E). We defined two sialylation gene signatures in melanocytes. The first one included the first five enzymes of the sialylation donor pathway (DP) (Fig. 1B). The second signature, termed sialylated glycoconjugates biosynthesis (SG bio), included the DP enzymes together with sialyltransferases (Fig. 1B). We found that malignant melanocytes had significantly higher expression of both signatures compared to healthy melanocytes (Fig. 1C). Using bulk RNA sequencing data from primary cutaneous melanoma patients from the The Cancer Genome Atlas (TCGA) we observed that the DP, but not the SG bio, was associated with lower progression-free survival (PFS) in these patients (Fig. 1D). Among the genes of the DP, we found that higher expression of *CMAS* correlated with lower PFS in patients with primary cutaneous melanoma (Fig. 1E, Supplementary Fig. S4F). CMAS catalyzes the conversion of *N*-acetylneuraminic acid (Neu5Ac, the most abundant sialic acid in humans) into the universal donor CMP-Neu5Ac required for all subsequent sialylation reactions (Fig. 1B). To investigate its role in tumor development, we knocked out *Cmas* in B16OVA murine melanoma cells, as we observed that cancer cells are the main cell type expressing high levels of sialic acids in subcutaneous B16OVA tumors (Supplementary Fig. S5A,B). *Cmas* deletion abolished surface sialic acids expression as determined by flow cytometry (Supplementary Fig. S5C). Using glycomics we confirmed a reduction in sialylated glycosphingolipids and *O*- and *N*-glycans from proteins in B16OVA *Cmas* KO cells compared to their MOCK counterparts, with an increase in non-sialylated structures on these molecules (Supplementary Fig. S5D,E; Supplementary Table S4A-C). Of note, knocking out *Cmas* did not affect cell viability *in vitro* (Supplementary Fig. S5F).

After subcutaneous injection in mice (Fig. 1F), the growth of *Cmas* KO tumors was heavily impaired compared to their MOCK counterparts (Fig. 1G). Tumor rejection was observed in 26 out of 38 *Cmas* KO-injected mice. In the remaining 12 mice, only small tumors developed (<100mm3) (Supplementary Fig. S5G), which contained higher infiltration of immune cells compared to MOCK tumors (Fig. 1H). The effect in tumor growth was not due to changes in the proliferation of tumor cells (Supplementary Fig. S5H) and B16OVA *Cmas* KO tumor cells maintained their sialic acid-negative phenotype *in vivo* (Supplementary Fig. S5I,J).

### *Cmas* deletion in tumor cells enhances anti-tumor lymphoid responses

To characterize the immune infiltrates of B16OVA MOCK and *Cmas* KO tumors, we performed spectral flow cytometry. We found an increased infiltration of CD4^+^ T cells and less regulatory T cells (Tregs) in B16OVA *Cmas* KO tumors compared to MOCK tumors, with no changes in CD8^+^ T cell infiltrates (Fig. 2A). Both CD4^+^ and CD8^+^ T cells infiltrating *Cmas* KO tumors presented an effector memory phenotype (Supplementary Fig. S6A,B) and expressed less inhibitory immune checkpoint molecules compared to T cells infiltrating MOCK tumors (Fig. 2B). Additionally, NK cells were increased in *Cmas* KO tumors and no changes were found in B cell infiltrations (Supplementary Fig. S6C,D).

**Figure 2.**
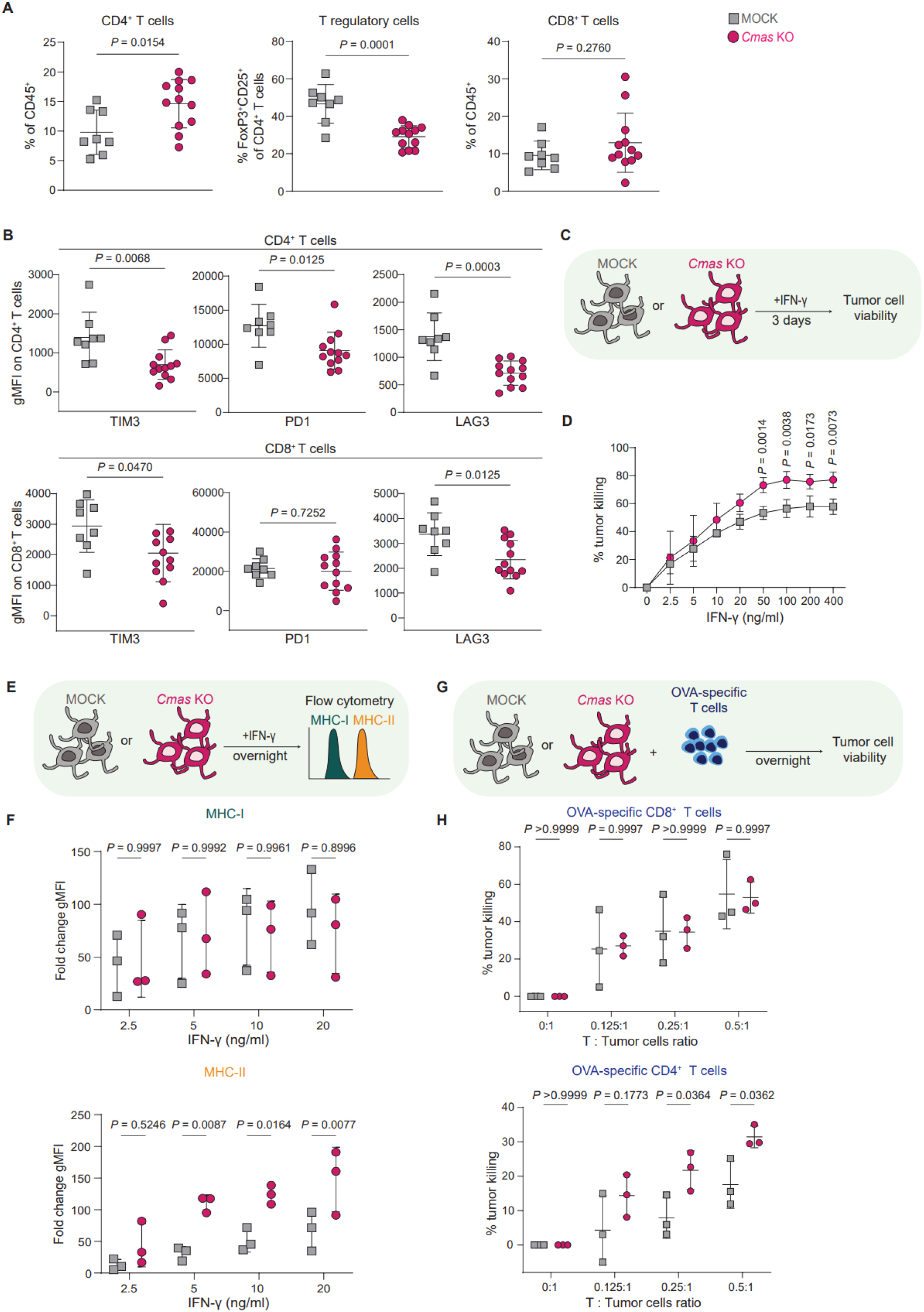
*Cmas* deletion in tumor cells enhances anti-tumor lymphoid responses. **A**, Frequency of different lymphoid cell subsets in B16OVA MOCK and *Cmas* KO tumors. Data shown as mean ± s.d. (n=8 mice in MOCK group and n=12 mice in *Cmas* KO group). Unpaired t-test used for statistical analysis. **B**, Expression of the immune checkpoints TIM3, PD1 and LAG3 on CD4^+^ (top) and CD8^+^ (bottom) T cells from B16OVA MOCK and *Cmas* KO tumors. Data shown as mean ± s.d. (n=8 mice in MOCK group and n=12 mice in *Cmas* KO group). Unpaired t-test used for statistical analysis. **C**, Scheme of IFN-γ-mediated killing experiment. B16OVA MOCK and *Cmas* KO tumor cells were cultured with increasing concentrations of IFN-γ for three days and cell viability was assessed using the CellTiter Blue assay. **D**, Percentage killing of B16OVA MOCK and *Cmas* KO cells after 3 days culture with IFN-γ. Data shown as mean ± s.d. from three independent experiments. Two-way ANOVA with Šídák’s multiple comparisons test used for statistical analysis. **E**, Scheme of the MHC induction experiment. B16OVA MOCK or *Cmas* KO were cultured overnight with increasing concentrations of IFN-γ and the expression of MHC-I and MHC-II molecules was assessed by flow cytometry. **F**, Expression of MHC molecules on B16OVA MOCK and *Cmas* KO cells upon stimulation with IFN-γ. Data from 3 independent experiments shown as mean ± s.d. of the fold change over the non-treated cells. Two-way ANOVA with Šídák’s multiple comparisons test used for statistical analysis. **G**, Scheme of T cell-mediated killing experiment. B16OVA MOCK and *Cmas* KO cells were co-cultured overnight with OVA-specific CD8^+^ and CD4^+^ T cells. After this, viability of the tumor cells was assed using CellTiter Blue assay and the percentage killing was calculated as described in the methods section. **H**, Percentage killing of B16OVA MOCK and *Cmas* KO cells by OVA specific CD8^+^ (top) and CD4^+^ (bottom) T cells. Data from 3 independent experiments. Nested one-way ANOVA with Šídák’s multiple comparisons test used for statistical analysis. gMFI: geometric Mean Fluorescence Intensity, s.d.: standard deviation, OVA: ovalbumin

We then asked whether genetic ablation of *Cmas* may induce tumor cell-intrinsic changes that can contribute to their susceptibility to lymphoid responses. First, we tested the susceptibility of tumor cells to interferon gamma (IFN-γ), as it induces apoptosis directly on cells and also regulates MHC molecules expression, impacting T cell activation. We observed increased killing of *Cmas* KO cells compared to MOCK cells upon IFN-γ treatment (Fig. 2C,D). Additionally, B16OVA *Cmas* KO cells had higher MHC-II expression than MOCK cells upon stimulation with IFN-γ, while MHC-I levels were increased similarly on both cell lines (Fig. 2E,F). Finally, adding to our previous results, we determined that B16OVA *Cmas* KO cells were more susceptible to OVA-specific CD4^+^ T cell-mediated killing than MOCK cells, while no differences were observed in OVA-specific CD8^+^ T cell-mediated killing (Fig. 2G,H). Altogether, deletion of *Cmas* in tumor cells improves the lymphoid landscape in murine melanoma and renders tumor cells intrinsically more susceptible to CD4^+^ T cell- and IFN-γ-mediated killing.

### *Cmas* KO tumor cells promote pro-inflammatory macrophage responses

As for the myeloid compartment, we found that *Cmas* KO tumors had a lower infiltration of Ly6C^high^ monocytes and higher infiltration of macrophages (Fig. 3A). Macrophages infiltrating *Cmas* KO tumors had increased expression of MHC-II and CD86, indicating a more pro-inflammatory phenotype compared to macrophages infiltrating MOCK tumors (Fig. 3B). We also found a significant increase in other myeloid cells infiltrations (Supplementary Fig. S7A,B), particularly in conventional dendritic cells type 2 (cDC2) (Supplementary Fig. S7C).

**Figure 3.**
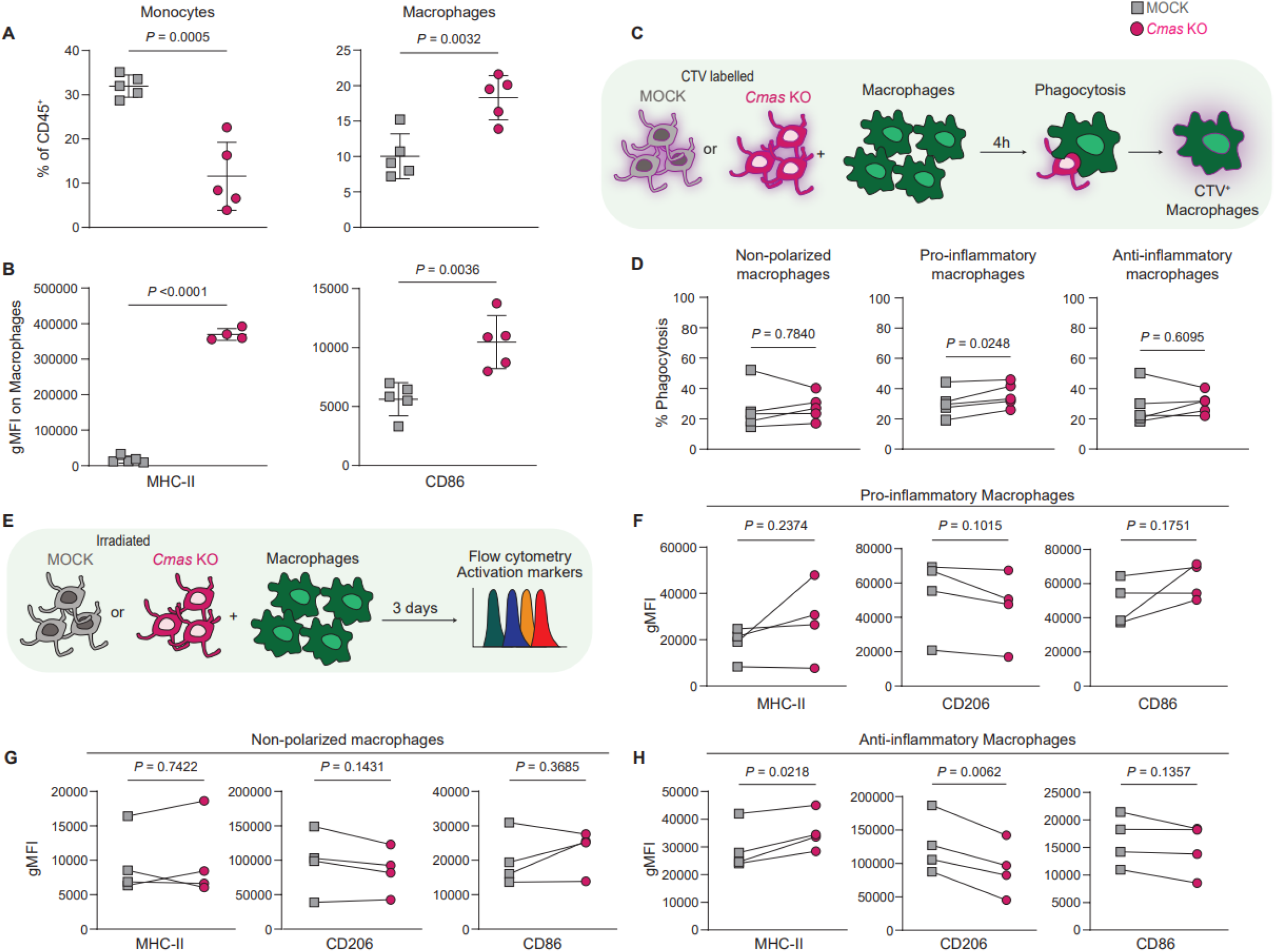
*Cmas* KO tumor cells promote pro-inflammatory macrophage responses. **A**, Frequency of monocytes (left) and macrophages (right) in B16OVA MOCK and *Cmas* KO tumors. Data shown as mean ± s.d. (n=5 mice per group). Unpaired t-test used for statistical analysis. **B**, Expression of MHC-II (left) and CD86 (right) on macrophages from B16OVA MOCK and *Cmas* KO subcutaneous tumors. Data shown as mean ± s.d. (n=5 mice per group). Unpaired t-test used for statistical analysis. **C**, Scheme of phagocytosis assay. B16OVA MOCK and *Cmas* KO cells were labelled with Cell Trace Violet (CTV) and co-cultured for four hours with either non-polarized, pro- or anti-inflammatory bone marrow-derived macrophages (BMDMs). After this, the percentage of macrophages that took up the CTV label (percentage phagocytosis) was assessed by flow cytometry. **D**, Percentage phagocytosis of B16OVA MOCK and *Cmas* KO cells by BMDMs. Data from 5 independent experiments. Paired t-test used for statistical analysis. **E**, Scheme of macrophage activation assay. Either non-polarized, pro- or anti-inflammatory BMDMs were co-cultured with B16OVA MOCK and *Cmas* KO cells for three days and the expression of different markers was determined using flow cytometry. **F-H**, Expression of MHC-II, CD86 and CD206 after co-culture with B16OVA MOCK and *Cmas* KO cells on pro-inflammatory (**F**), non-polarized (**G**) and anti-inflammatory (**H**) macrophages. Data from 4 independent experiments. Unpaired t-test used for statistical analysis. gMFI: geometric Mean Fluorescence Intensity, s.d.: standard deviation, OVA: ovalbumin

As we found an increase in pro-inflammatory macrophages in *Cmas* KO tumors *in vivo*, we investigated whether *Cmas* deletion increases tumor cell-susceptibility to macrophage phagocytosis, leading to a more efficient activation. We observed increased phagocytosis of *Cmas* KO cells by pro-inflammatory macrophages compared to B16OVA MOCK cells (Fig. 3C,D). However, after a longer co-culture of *Cmas* KO and MOCK B16OVA cells with pro-inflammatory macrophages, we did not observe significant differences in the expression of CD86, CD206 or MHC-II on them (Fig. 3E,F). Thus, we asked whether B16OVA *Cmas* KO tumor cells promoted a pro-inflammatory phenotype on non-polarized or anti-inflammatory macrophages. We did not observe changes in CD86, CD206, or MHC-II expression on non-polarized macrophages, however, we found a significant increase in MHC-II and decrease in CD206 expression on anti-inflammatory macrophages when co-cultured with *Cmas* KO cells (Fig. 3G,H). These findings indicate that *Cmas* KO tumor cells promote infiltration and activation of macrophages in the TME and are not only more susceptible to phagocytosis, but also induce pro-inflammatory phenotypic changes in anti-inflammatory macrophages.

### Tumor cells with low *CMAS* expression are proximal to T cells and macrophages in human melanoma

Our murine model demonstrated that *Cmas*-deficient melanoma cells promote T cell and macrophage responses. However, in humans there are no *CMAS*-deficient cancers, instead *CMAS* is a broadly expressed gene. To explore this heterogeneity *in situ* and its relationship with the immune TME in human melanoma, we performed single-cell spatial transcriptomics. We analysed a cohort of 4 neoadjuvant ICB-treated patients (2 responders and 2 non-responders) with resectable stage III melanoma and matched pre- and post-ICB treatment samples (Fig. 4A), using a customized gene panel that included immune-oncology reference genes and multiple sialylation–related enzymes, including *CMAS*. Data from seven tissue sections were included (Fig. 4B, Supplementary Fig. S8), with no consistent differences in immune cell infiltration across samples (Supplementary Fig. S9A).

**Figure 4.**
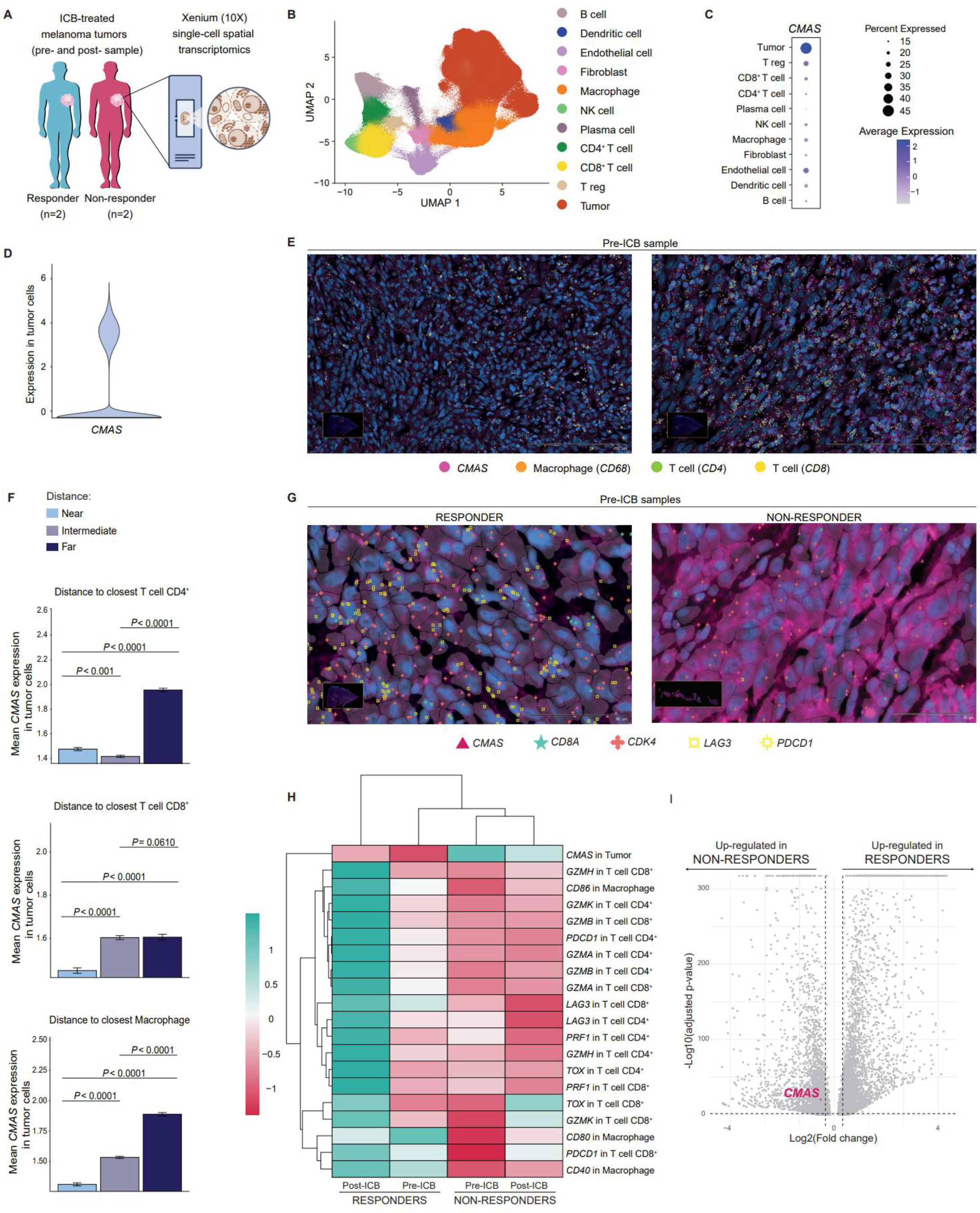
*CMAS* expression in tumor cells inversely correlates with distance to and activation of immune cells, and with ICB response in human melanoma. **A**, Scheme of spatial transcriptomics ICB-treated melanoma patient cohort (n=4), three ICB-treated melanoma patient samples pre- and post-ICB, and one patient only pre-ICB were included, in a total of seven tumor tissue slides. **B**, UMAP plot of all cells from tissues described in (A). Indicated the main cell types annotated in the dataset. **C**, Percentage and average expression of CMAS across cell types identified in (B). **D**, Violin plot of CMAS expression in tumor cells in all pre-ICB treatment samples. **E**, Example pictures of a pre-ICB patient sample (partial responder). Dots depict expression of genes: CMAS (pink), CD68 (orange, macrophage marker), CD4 (green, T cell marker) and CD8 (yellow, T cell marker). Size of the dot indicates expression abundance. **F**, Mean CMAS expression in tumor cells relative to distance groups from the closest target cell. “Near” corresponds to the first tercile, “Intermediate” to the second tercile, and “Far” to the third tercile of all distances calculated. Bar plots show the average CMAS expression per group with standard deviation. Wilcoxon test with Bonferroni correction used for statistical analysis. Analysis performed on pre-ICB samples. **G**, Example pictures of melanoma responders and non-responders TME immune composition pre-ICB. Icons depict expression of genes: CMAS (pink triangle), CDK4 (red cross, tumor marker), CD8 (blue star, T cell marker), LAG3 (yellow circle, immune checkpoint) and PDCD1 (yellow square, immune checkpoint). Size of the icon indicates expression abundance. **H**, Heatmap showing gene expression of the indicated gene in the indicated cell type (eg. CMAS gene expression in tumor cells). Unsupervised clustered. **I**, Differential gene expression analysis on tumor cells of four ICB-treated melanoma patients from an independent single-cell RNA sequencing cohort (two responders and two non-responders). Dotted lines indicate P<0.05 and log2(fold change)>0.5. TME: tumor microenvironment, ICB: immune checkpoint blockade, T reg: regulatory T cell, NK: natural killer

First, we evaluated *CMAS* expression in all cell types before ICB treatment and found that *CMAS* was predominantly and heterogeneously expressed in tumor cells (Fig. 4C,D). In addition, we also observed a heterogeneous distribution of immune cells (such as T cells and macrophages), with some areas being more infiltrated than others in the same tissue sample (Fig. 4E). Therefore, we asked whether *CMAS* expression in tumor cells was associated with these immune distributions. However, we did not observe clear areas of *CMAS* high and low expression, instead *CMAS* low and high-expressing tumor cells were distributed across all the tissue (Fig. 4E). We then calculated for each individual tumor cell in pre-ICB samples its distance to the closest CD4^+^, CD8^+^ T cell or macrophage. We observed that tumor cells in closer proximity to T cells and macrophages expressed lower *CMAS* (Fig. 4F). We performed this analysis for all cell types identified (Fig. 4B), finding distinct trends per cell type (Supplementary Fig. S9B). We also analysed post-treatment samples, where the heterogeneous expression of *CMAS* in tumor cells and its correlation with distance to T cells and macrophages showed the same results (Supplementary Fig. S9C,D). Altogether these data show that the heterogeneous distribution of *CMAS*-expressing tumor cells and immune cells in the TME is non-stochastic, being tumor cells with lower *CMAS* expression closer to defined immune cell types.

### *CMAS* expression in tumor cells inversely correlates with immune activation and ICB response in melanoma patients

Finally, we assessed whether CMAS is associated with the activation status of these immune cell subsets. We determined *CMAS* expression in tumor cells in the different patients (responders or non-responders) pre- and post-ICB treatment, as well as the expression of defined markers for activation of macrophages and T cells and immune checkpoints (Fig. 4G). Using unsupervised clustering we observed that these immune markers were lower expressed in non-responders compared to responders, both before and after ICB treatment (Fig. 4H). The same trends were observed for activation markers on DCs and NK cells (Supplementary Fig. S10A). In contrast, *CMAS* expression in tumor cells followed an inversed pattern, being higher expressed in non-responders compared to responders, both pre- and post-treatment (Fig. 4H). We validated this finding in an independent single-cell sequenced neoadjuvant ICB-treated melanoma patient cohort, finding *CMAS* within the significantly upregulated genes in non-responders (Fig. 4I, Supplementary Fig. S10B,C).

## Discussion

We here show that expression of the gene *CMAS* (which encodes the enzyme responsible for activating sialic acid as the essential donor for sialylation) in tumor cells correlates with poor response to immunotherapy in melanoma patients. Using spatially resolved analysis, we found that higher *CMAS* expression in tumor cells is associated with increased physical distance between tumor cells and both T cells and macrophages in the tumor microenvironment, linking for the first time the expression of tumor-sialylation genes with the spatial organization of tumors. In a murine melanoma model, we confirmed that deletion of *Cmas* abrogates tumor growth, dampening the immune tolerance imposed by sialic acids on both lymphoid and myeloid cells. In particular, *Cmas* deletion increased tumor cell-susceptibility to CD4^+^ T cell- and IFN-γ-mediated killing, as well as phagocytosis and the inflammatory state of macrophages.

Aberrant sialylation is a common feature across tumor types, largely driven by altered expression of sialylation pathway enzymes in cancer cells ^6,14^. Our findings align with prior reports in which *Cmas* deletion delayed breast cancer growth ^12^. However, in murine pancreatic cancer, deleting *Cmas* had no impact in tumor growth, but when combined with immune checkpoint blockade, enhanced anti-tumor immunity was observed ^11^, indicating that even in immune-cold tumors sialylation may improve therapeutic potential of ICB ^9^. Whether the expression of the *CMAS* gene may also correlate with survival and response to therapy in these tumor types remains to be determined.

Although deletion of genes of the donor pathway, such as *Gne* and *Slc35a1*, delayed tumor growth in murine melanoma, the effect on these models was not as strong as we report here with *Cmas* KO, which resulted in a complete lack of sialic acids ^9,32^. Even though *CMAS* deletion cannot be a therapeutic option, other strategies that aim to reduce sialic acids have shown to increase anti-tumor immunity, such as a metabolic inhibitor of sialyltransferases or targeted sialidases ^9,10^. However, in these cases sialylation was reduced not only on tumor cells but also on other cells in the TME, such as immune cells and stromal cells, which also influence immune responses ^33-35^. Recently, it was found that CMAS expression is regulated by micro RNAs, which directly impact surface sialylation ^36^. Further research is needed to determine whether these miRNAs affect CMAS expression in the TME, and if they may be useful as treatments and therefore change patient outcome.

We observed that *Cmas* KO melanoma tumors had increased infiltration of less exhausted T cells and pro-inflammatory macrophages. Similar results in the immune TME were seen with either genetic or therapeutic reduction of sialylation in other murine tumor models ^9-12,37^. Sialic acids are known to interact with inhibitory sialic acid-binding receptors expressed on immune cells, called Siglecs (Sialic acid-binding immunoglobulin like lectins), which dampen lymphoid and myeloid cell responses ^6,38^. Although the lack of surface sialylation on *Cmas* KO tumor cells may reduce inhibitory signals through Siglecs, it also exposes many non-sialylated glycan structures, as we defined using glycomics, which can bind other glycan-binding receptors on immune cells impacting their activation. In addition, sialylated glycoconjugates play important tumor cell-intrinsic roles that can affect tumor development independently from Siglecs, such as regulation of receptors activation, migration and proliferation ^14,39-42^. In line with this, we observed that B16OVA *Cmas* KO tumor cells are intrinsically more susceptible to IFN-γ-mediated killing. Moreover, IFN-γ caused a higher upregulation of MHC-II molecules on *Cmas* KO tumor cells compared to MOCK cells. In agreement with our findings, increased expression of MHC molecules was seen on dendritic cells upon desialylation and on *Cmas* KO breast cancer cells ^12,35^. This means that *Cmas* KO tumor cells may be intrinsically more capable of interacting and presenting antigens to CD4^+^ T cells, which makes them more susceptible to CD4^+^ T cell-mediated killing, as we observed. Perdicchio et al. determined that the decrease in tumor growth in the B16OVA *Slc35a1* KO model was dependent on NK cells, which promoted infiltration of CD4^+^ T cells and IFN-γ release ^32^. In line with this study, we found an increase not only in NK cells, but also in MHC-II^high^ macrophages and cDC2s infiltration in *Cmas* KO tumors, which may also favour CD4^+^ T cell responses.

Using single-cell spatial transcriptomics, we demonstrated that *CMAS* is heterogeneously expressed in tumor cells of melanoma patients, and tumor cells that have lower *CMAS* expression are more proximal to T cells, macrophages, and other immune cell types in the TME. One possible explanation is that infiltrating immune cells find a favourable niche close to *CMAS*^low^ tumor cells because they are more capable of interacting with them; and *CMAS*^low^ tumor cells are intrinsically more susceptible to killing, as we showed in our murine model. Previous work has shown that the spatial arrangement of T cells and macrophages in the TME impact patient survival and response to ICB therapy in melanoma ^17,18,43,44^. We found that tumor *CMAS* expression is increased in non-responders to neoadjuvant ICB therapy, being inversely correlated with immune activation and proximity of immune cells to tumor cells. Interestingly, we also observed that macrophages are more activated in responder patients than in non-responders before ICB therapy, suggesting that the combination of *CMAS* expression in tumor cells and macrophages activation may be a good predictor of ICB response. Due to the limited size of our cohorts of patients, the value of CMAS as a predictor of patient response to ICB needs to be confirmed in more extensive cohorts.

Altogether, our results position CMAS as a key immune regulator and as a promising predictive marker of patient outcome and response to neoadjuvant ICB in melanoma.

## Acknowledgements

We acknowledge Sara Rohani from the Amsterdam UMC, location AMC, for her help with the patient samples for spatial transcriptomics. MC would like to acknowledge Amber Genser, Vinicio Melo, Andreia Linhares Miranda for their help in *in vivo* experiments, Sandra van Vliet for her insightful feedback, and all the melanoma patients who accepted to donate their samples to research. We also acknowledge our funding resources: European Union Horizon 2020, Marie Skłodowska-Curie Actions Grant agreement No. 956758, GLYTUNES Consortium (MC), SPINOZA NWO SPI-93-538 (SIM, BS, ER), Spatial Biology Proof of Concept grant from the Microscopy and Cytometry Core facility from the Amsterdam UMC, location VUmc (SIM, MC), Dutch Cancer Foundation KWF16684 (SW), Oncode Accelerator, a Dutch National Growth Fund project under grant number NGFOP2201 (GC), European Union Horizon 2020, Marie Skłodowska-Curie Actions Grant agreement No. 859974, EDUC8 Consortium (EN), The HealthHolland program TARGlycan (TKI-LSH-DT2O22LUMC:2022-02) and the EU HORIZON-CSA program GLYCOTwinning (101079417) (TZ, NdH)

## Notes

### Competing Interest Statement

The authors have declared no competing interest.

