## Supplementary Figures and Tables for "CMAS dampens anti-tumor immunity and associates with response to neoadjuvant immunotherapy in melanoma"

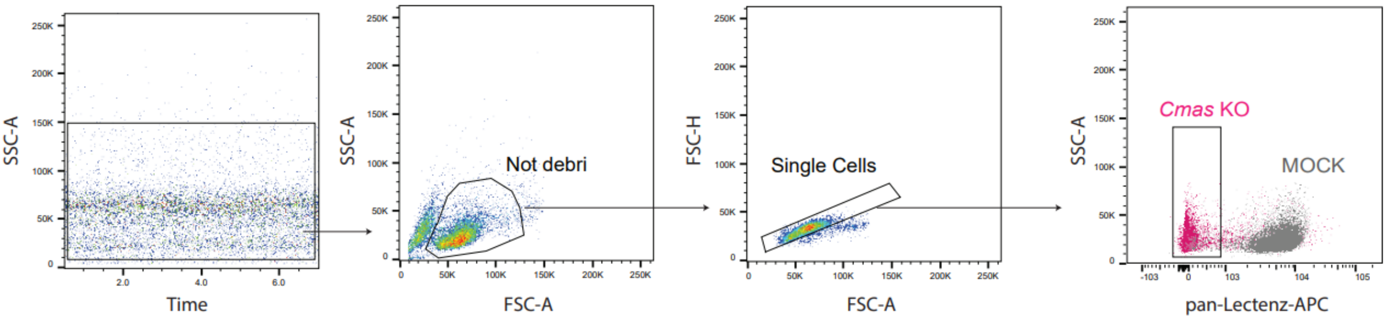


**Supplementary Figure S1. Gating strategy for the sorting of B16OVA *Cmas* KO cells.** After removing unstable events over time, debri and doublets, *Cmas* KO tumor cells were sorted based on negative pan-Lectenz staining.

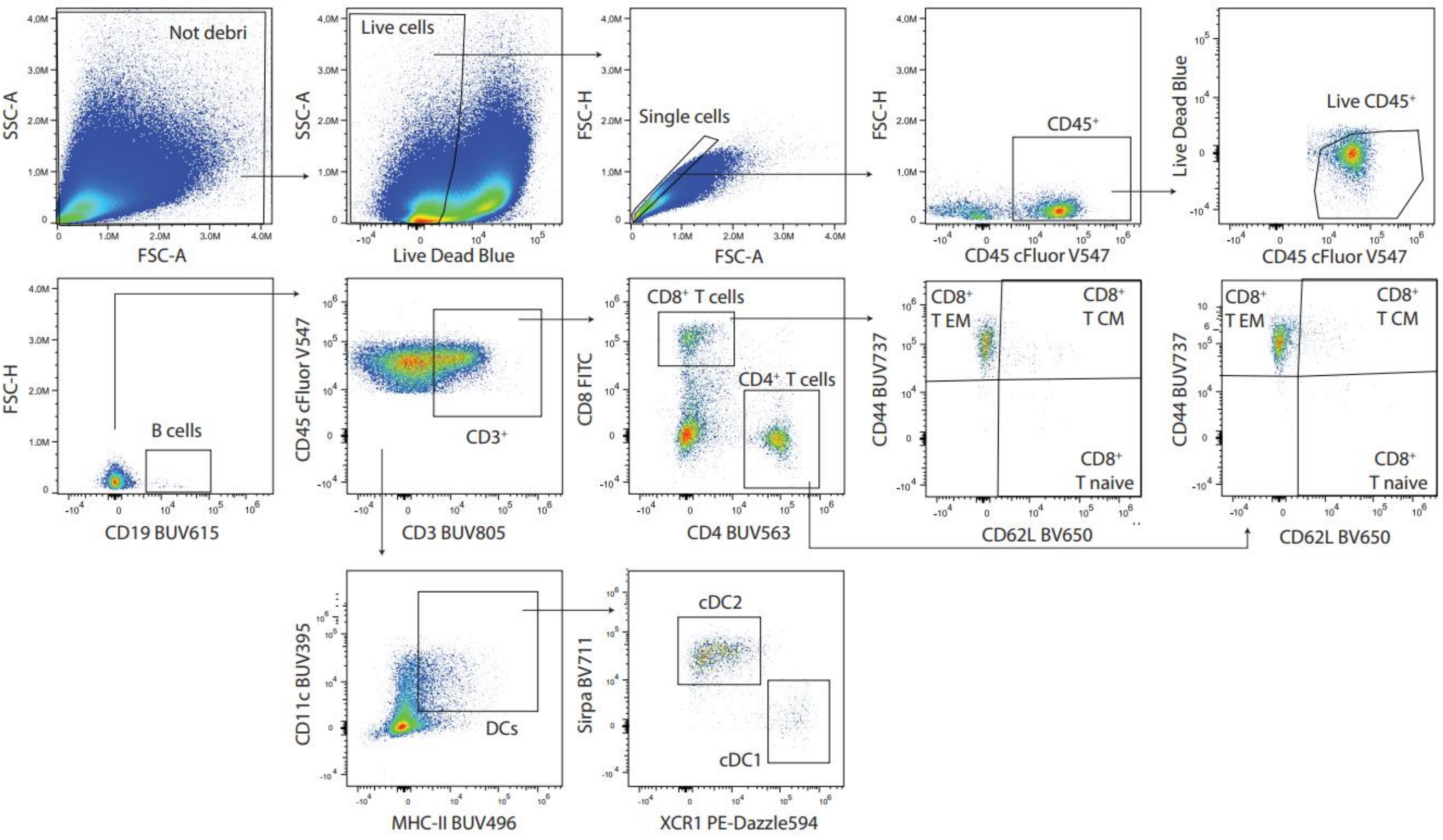


**Supplementary Figure S2. Gating strategy for panel 1.** Gating strategy used for immune cell subsets defined using panel 1. Antibodies can be found in Supplementary Table S2. EM: effector memory, CM: central memory, DCs: dendritic cells, cDC: conventional dendritic cell

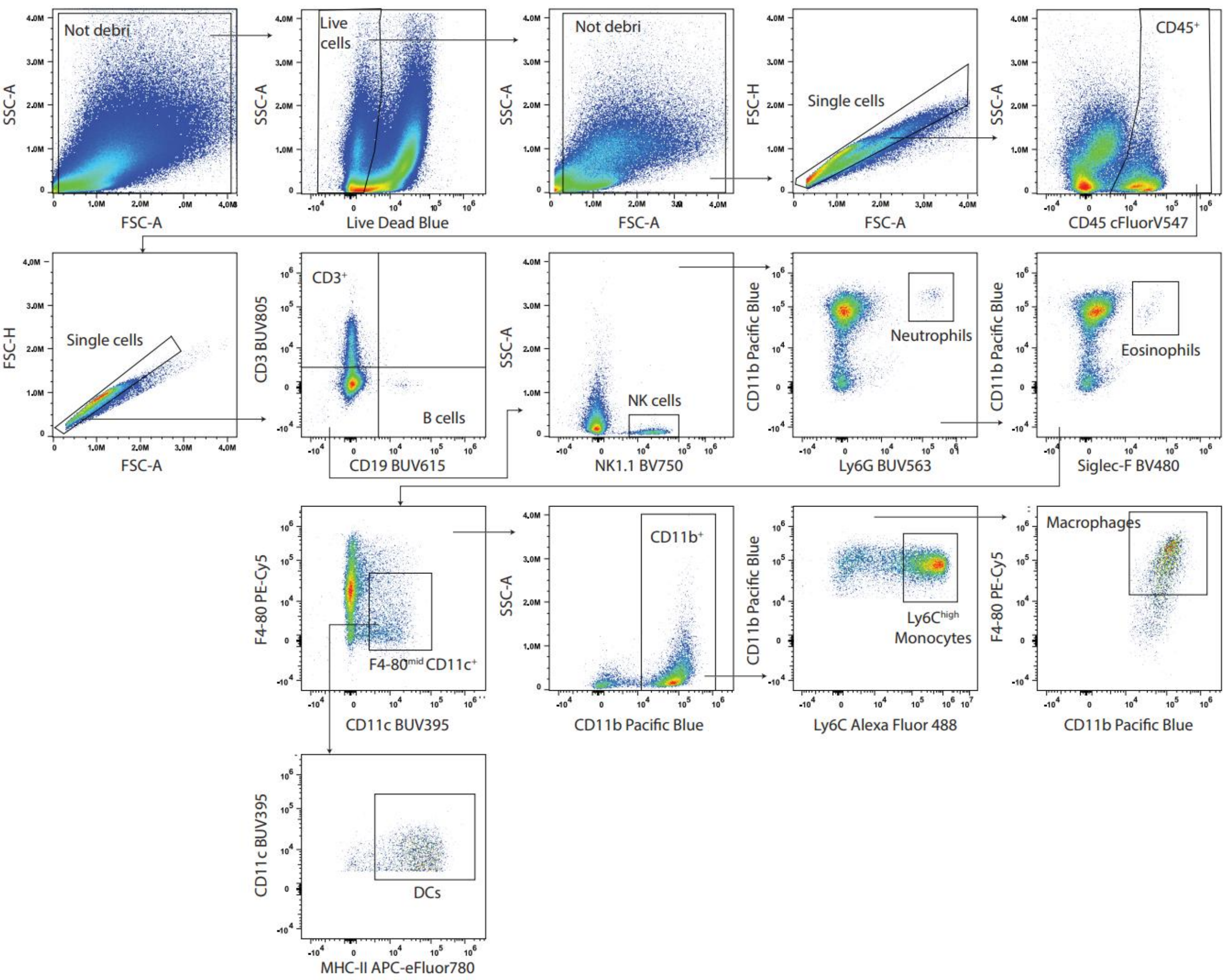


**Supplementary Figure S3. Gating strategy for panel 2.** Gating strategy used for immune cell subsets defined using panel 2. Antibodies can be found in Supplementary Table S2. NK: natural killer, DCs: dendritic cells.

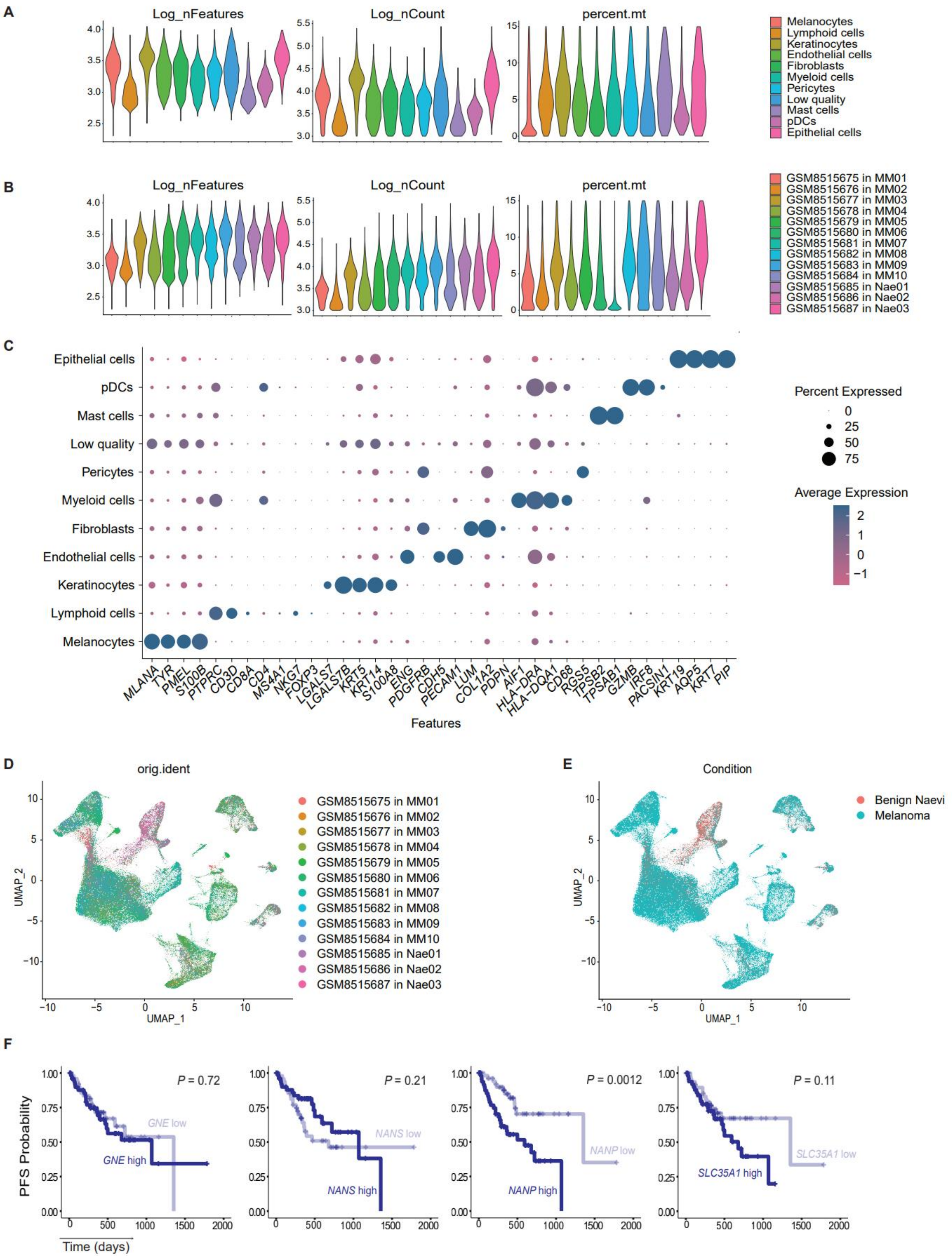


**Supplementary Figure S4. Analysis of scRNA-seq and survival data from TCGA.**

**A,B,** Violin plots showing the logarithm of nFeatures (number of genes expressed per cell), nCounts (total transcripts count) and percent.mt (percentage of mitochondrial genes expression relative to total gene expression) after filtering, per cluster (**A**) or per sample (**B**). **C,** Dot plot showing the expression of features (genes) in the 11 clusters defined. Dot size represents the percentage of cells expressing the gene, and the colour is based on the average expression level. **D,E,** UMAP plots coloured by sample (**D**) or condition (benign naevi or melanoma)(**E**). **F,** Kaplan-Meier plots of progression-free survival (PFS) in primary melanoma patients from TCGA with low (n=52) or high (n=51) expression, based on the median, of enzymes from the donor pathway of sialylation. Statistical test: Log-rank test.

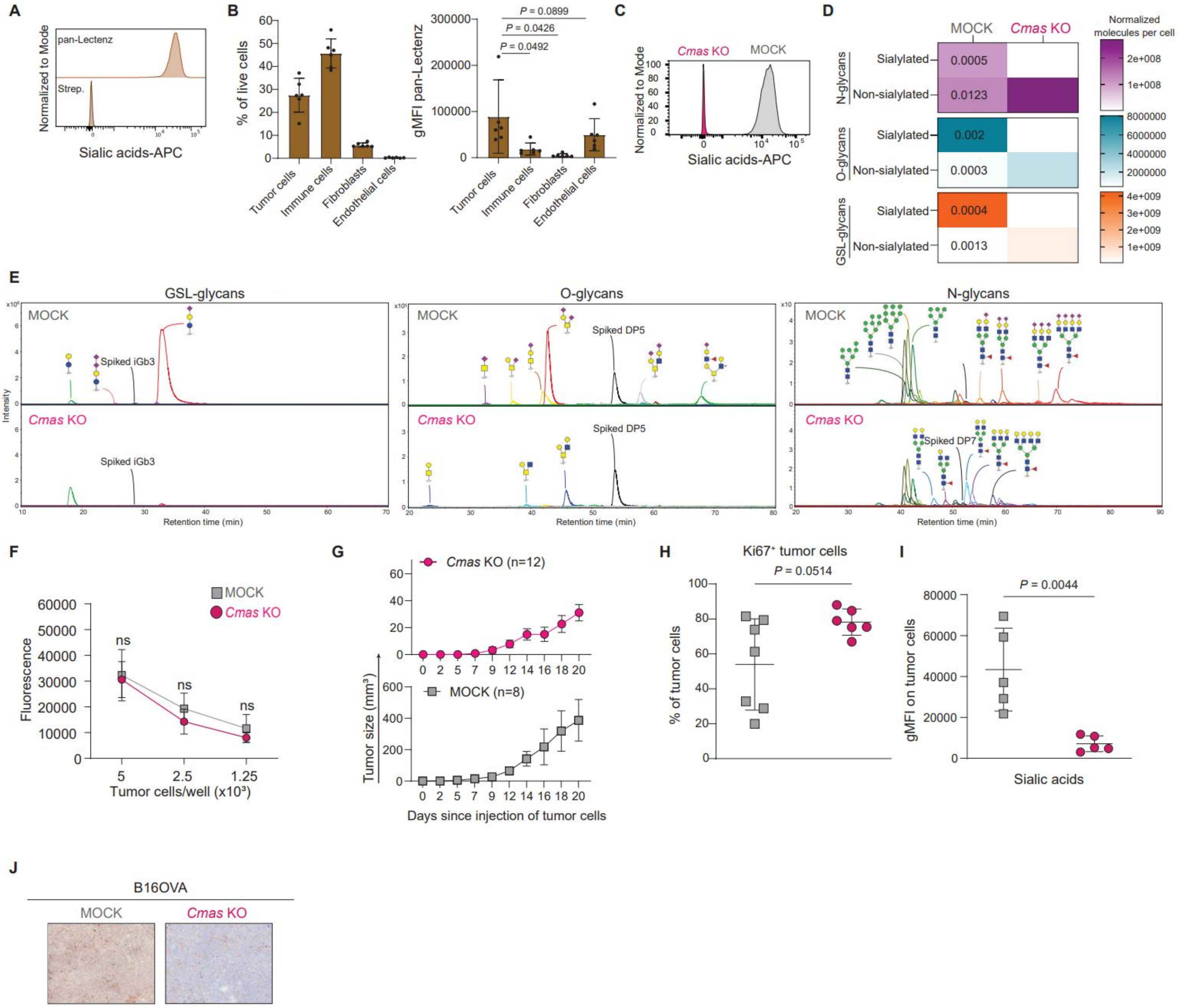


**Supplementary Figure S5. B16OVA *Cmas* KO *in vivo* model. A,** Histograms of sialic acids expression on B16OVA wild type (WT) cells determined by flow cytometry. Strep.= streptavidin only negative control. **B,** Bar plots showing the frequency of four major cell types in B16OVA WT subcutaneous tumors (left) and the expression of sialic acids on those cell types (right). Statistical analysis: paired t-test between each cell type and tumor cells. **C,** Histograms of sialic acids expression on B16OVA MOCK and *Cmas* KO cells determined by flow cytometry. **D,** Heatmaps showing the normalized abundance of sialylated and non-sialylated N-, O-glycans and glycosphingolipid (GSL)-glycans per cell in B16OVA MOCK and *Cmas* KO cells determined using mass spectrometry-based glycomics. Data is from three technical replicates. Statistical analysis: unpaired t-test. **E,** Profiles of GSL-glycans, O-glycans and N-glycans found on B16OVA MOCK and *Cmas* KO cells determined by mass spectrometry. **F,** Viability of B16OVA MOCK and *Cmas* KO cells using the CellTiter Blue assay. Data from three independent experiments. Data shown as mean ± s.d. Statistical analysis: Two-way ANOVA with with Šídák's multiple comparisons test. **G,** Size of B16OVA MOCK and *Cmas* KO subcutaneous tumors used for the analysis of the tumor immune microenvironment. Data shown as mean ± s.e.m. **H,** Percentage of proliferative tumor cells (Ki67+) from B16OVA MOCK and *Cmas* KO tumors. Data shown as mean ± s.d. Statistical analysis: unpaired t-test. **I,** Expression of sialic acids on tumor cells from B16OVA MOCK and *Cmas* KO subcutaneous tumors. Data shown as mean ± s.d. Statistical analysis: unpaired t-test. **J,** Immunohistochemistry staining of sialic acids on B16OVA MOCK and *Cmas* KO subcutaneous tumors. Sialic acid signal is depicted in brown. gMFI: geometric Mean Fluorescence Intensity

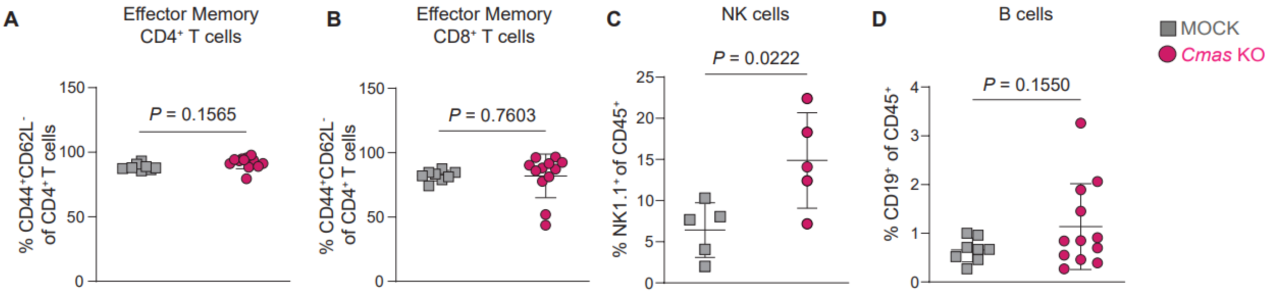


**Supplementary Figure S6. Infiltration of lymphoid cells in MOCK and *Cmas* KO tumors. A,B,** Frequency of effector memory CD4+ (**A**) and CD8+ (**B**) T cells in B16OVA MOCK (n=8) and *Cmas* KO (n=12) subcutaneous tumors. **C,** Percentage of Natural Killer (NK) cells in B16OVA MOCK and *Cmas* KO subcutaneous tumors (n=5 mice per group). **D,** Percentage of B cells in B16OVA MOCK (n=8) and *Cmas* KO (n=12) subcutaneous tumors. Data shown as mean ± s.d. Statistical analysis: unpaired t-test. gMFI: geometric Mean Fluorescence Intensity.

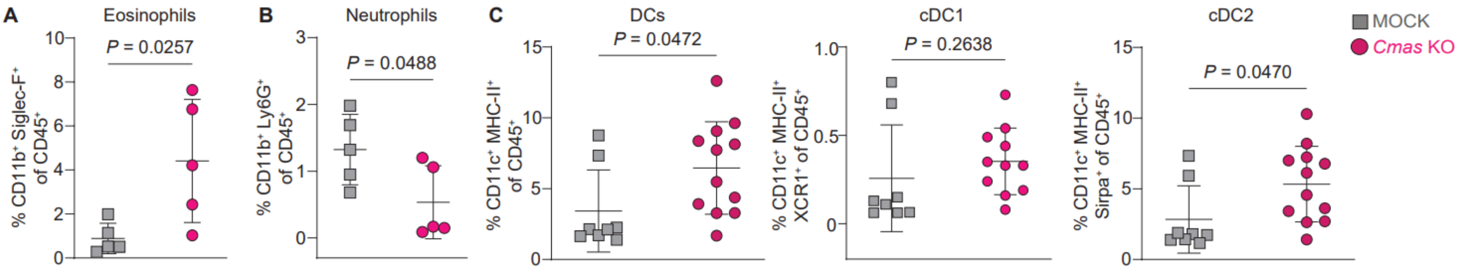


**Supplementary Figure S7. Infiltration of myeloid cells in MOCK and *Cmas* KO tumors. A-C,** Percentage of eosinophils (**A**), neutrophils (**B**), dendritic cells (**DCs**), conventional DCs (cDC) 1 and cDC2 (**C**) in B16OVA MOCK and *Cmas* KO subcutaneous tumors. Data shown as mean ± s.d. (n=8 mice in MOCK group and n=12 in *Cmas* KO group). Statistical analysis: unpaired t-test.

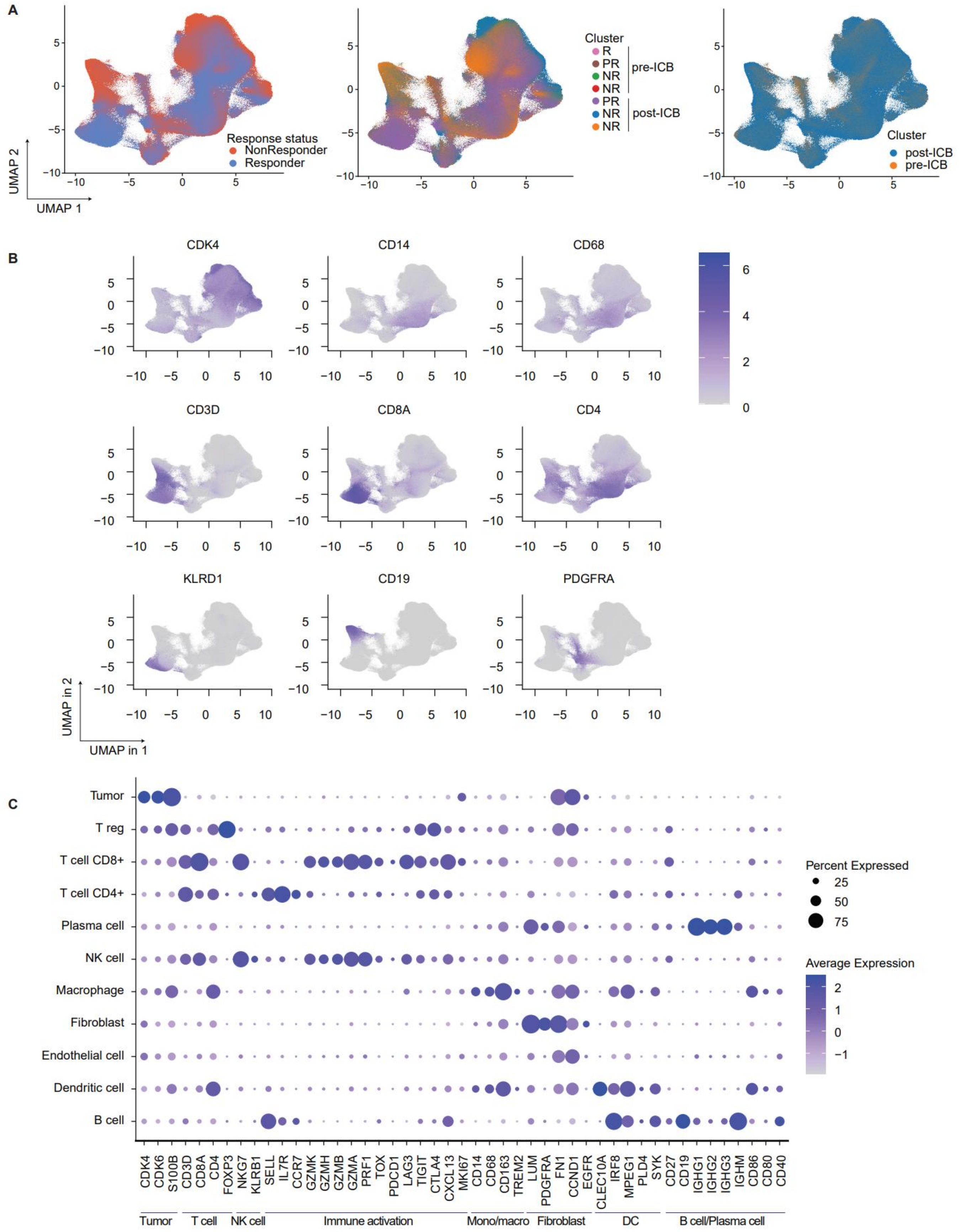


**Supplementary Figure S8. Analysis of single-cell spatial transcriptomics. A,**

UMAP of all cells from Fig. 4B. Cells are grouped by response (left), patient (center) and treatment timepoint (right). **B,** UMAP of all cells from Fig. 4B. Expression of main lineage markers is shown. Tumor marker (*CDK4*), monocyte/ macrophages markers (*CD14, CD68*), T cell markers (*CD3D, CD8A and CD4*), NK cell marker (*KLRD1*), B cell marker (*CD19*) and fibroblast marker (*PDGFRA*). **C,** Percentage and average expression of lineage and functional markers per annotated cell cluster. ICB: immune checkpoint blockade, NK: Natural killer, DC: Dendritic cell, R: Responder, PR: Partial responder, NR: Non-responder.

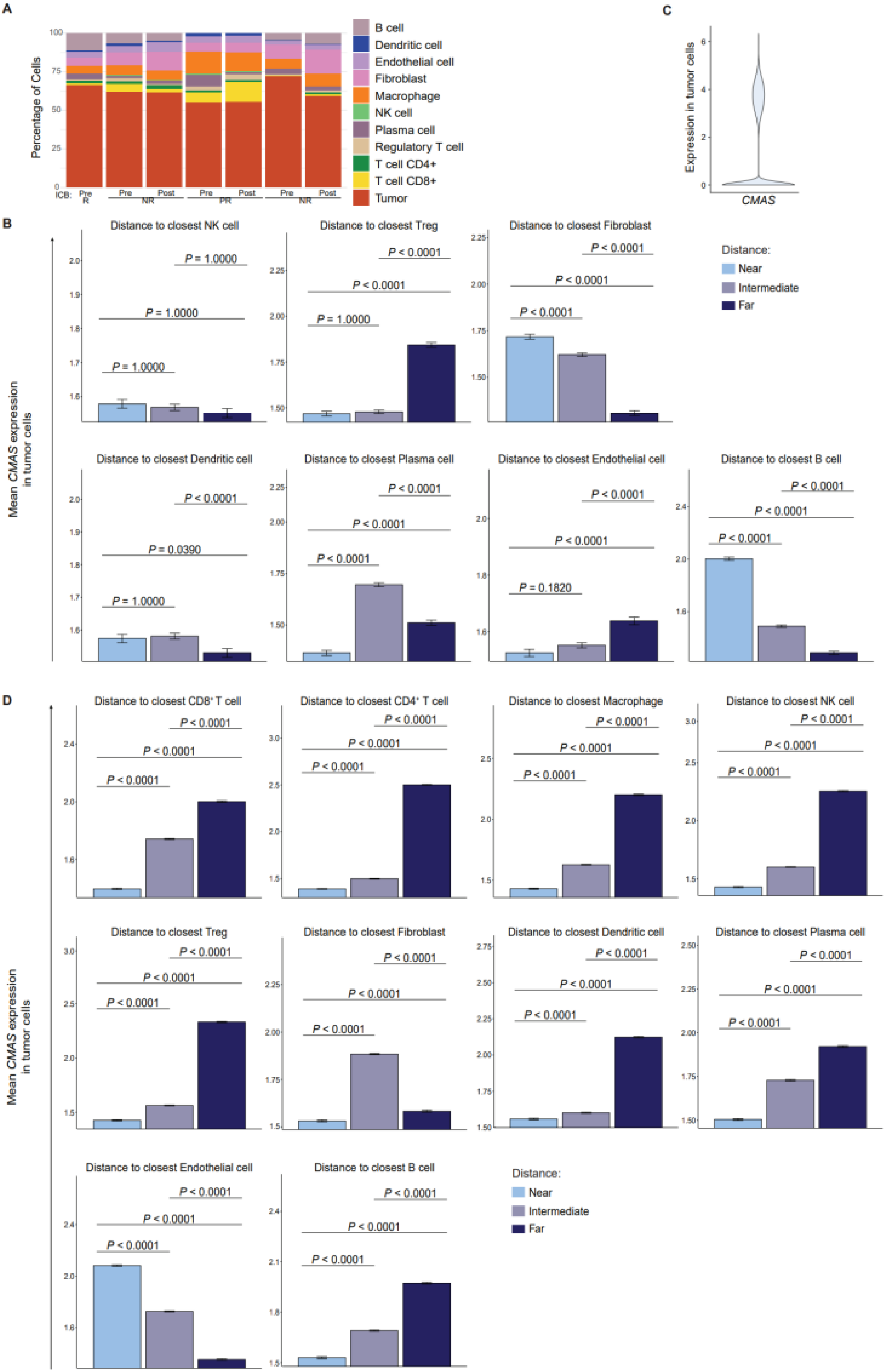
A
Percentage of Cells
ICB: Pre R
Pre Post NR
Pre Post PR
Pre Post NR
B cell
Dendritic cell
Endothelial cell
Fibroblast
Macrophage
NK cell
Plasma cell
Regulatory T cell
T cell CD4+
T cell CD8+
Tumor
C
Expression in tumor cells
CMAS
B
Mean CMAS expression in tumor cells
Distance to closest NK cell
P = 1.0000
P = 1.0000
P = 1.0000
Distance to closest Treg
P < 0.0001
P < 0.0001
P = 1.0000
Distance to closest Fibroblast
P < 0.0001
P < 0.0001
P < 0.0001
Distance:
Near
Intermediate
Far
Distance to closest Dendritic cell
P < 0.0001
P = 0.0390
P = 1.0000
Distance to closest Plasma cell
P < 0.0001
P < 0.0001
P < 0.0001
Distance to closest Endothelial cell
P < 0.0001
P < 0.0001
P = 0.1820
Distance to closest B cell
P < 0.0001
P < 0.0001
P < 0.0001
D
Mean CMAS expression in tumor cells
Distance to closest CD8+ T cell
Distance to closest CD4+ T cell
Distance to closest Macrophage
Distance to closest NK cell
Distance to closest Treg
Distance to closest Fibroblast
Distance to closest Dendritic cell
Distance to closest Plasma cell
Distance to closest Endothelial cell
Distance to closest B cell
P < 0.0001
Distance:
Near
Intermediate
Far

**Supplementary Figure S9. Spatial transcriptomics on melanoma patient tissues.** **A,** Average frequencies of annotated cell clusters in Fig. 4B across different patient tissues. ICB= immune checkpoint blockade, R= responder, NR= non-responder. **B,** Mean *CMAS* expression in tumor cells relative to distance groups from the nearest target cell. “Near” corresponds to the first tercile, “Intermediate” to the second tercile, and “Far” to the third tercile of all the distances calculated. Bar plots show the average *CMAS* expression per group with standard deviation. Statistical significance was assessed using the Wilcoxon test with Bonferroni correction. Analysis performed on pre-ICB samples. **C,** Violin plot of *CMAS* expression in tumor cells in all post-ICB treatment samples. **D,** Mean *CMAS* expression in tumor cells relative to distance groups from the nearest target cell. “Near” corresponds to the first tercile, “Intermediate” to the second tercile, and “Far” to the third tercile of all the distances calculated. Bar plots show the average *CMAS* expression per group with standard deviation. Statistical significance was assessed using the Wilcoxon test with Bonferroni correction. Analysis performed on post-ICB samples. NK: natural killer, Treg: regulatory T cells, R: Responder, PR: Partial responder, NR: Non-responder

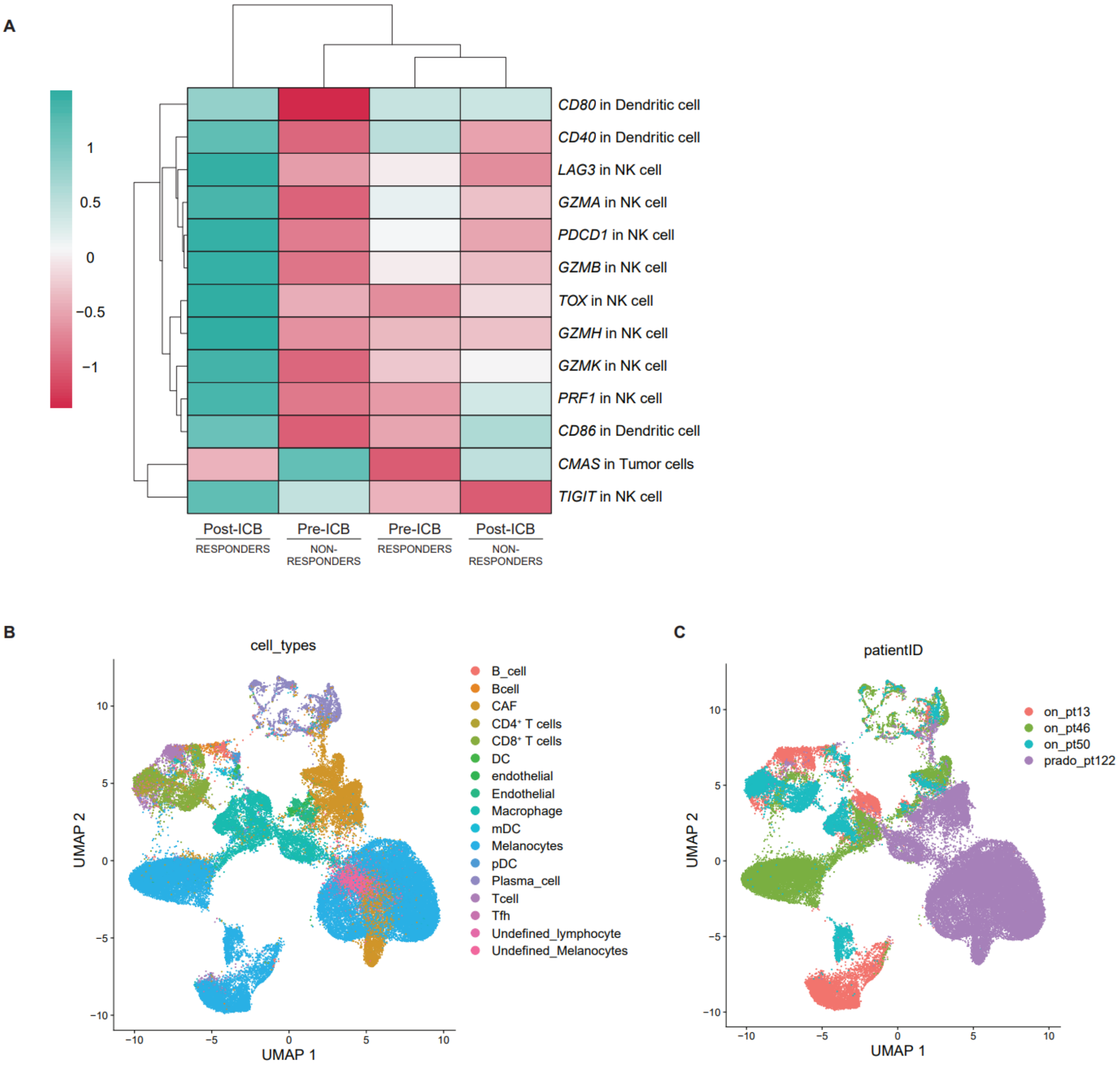


**Supplementary Figure S10. *CMAS* expression in tumor cells correlates with immune activation and ICB response. A,** Heatmap showing gene expression of the indicated gene in the indicated cell type (eg. *CMAS* gene expression in tumor cells). Unsupervised clustered. **B,C,** UMAP plots of scRNA-seq data from Fig. 4I, coloured by cell types (**B**) or patient ID (**C**). ICB: immune checkpoint blockade, NK= natural killer, DC: dendritic cell

### Supplementary Tables

**Supplementary Table S1. Antibodies included in panels 1 and 2 used for immune profiling of murine B16OVA MOCK and Cmas KO tumors by spectral flow cytometry**. List of antibodies used for staining for spectral flow cytometry of murine tumors. Those included in panel 1 and/or panel 2 are marked with a cross.

| Antibody | Fluorochrome | Company | Catalogue # | Panel 1 | Panel 2 |
|---|---|---|---|---|---|
| CD8b Monoclonal Antibody (eBioH35-17.2 (H35-17.2)) | FITC | Invitrogen | 11-0083-82 | X | |
| BD OptiBuild™ Hamster Anti-Mouse CD3 | BUV805 | Becton Dickinson | 749276 | X | X |
| BD Horizon™ Rat Anti-Mouse CD4 | BUV563 | Becton Dickinson | 612923 | X | |
| Anti-mouse CD279 (PD-1) Antibody | BV785 | Biolegend | 135225 | X | |
| BD Horizon™ Rat Anti-Mouse CD44 | BUV737 | Becton Dickinson | 612799 | X | |
| Anti-mouse CD62L Antibody | BV650 | Biolegend | 104453 | X | |
| Anti-Mouse CD45 (30-F11) | cFluorV547 | Cytek | SKU R7-20571 | X | X |
| BD Horizon™ Hamster Anti-Mouse CD11c | BUV395 | Becton Dickinson | 564080 | X | X |
| Anti-mouse/human CD11b Antibody | Pacific Blue | Biolegend | 101224 | X | X |
| BD OptiBuild™ Rat Anti-Mouse I-A/I-E (MHC-II) | BUV496 | Becton Dickinson | 750281 | X | |
| Anti-mouse/rat XCR1 Antibody | PE-Dazzle594 | Biolegend | 148234 | X | X |
| BD OptiBuild™ Rat Anti-Mouse CD172a | BV711 | Becton Dickinson | 740766 | X | X |
| BD OptiBuild™ Rat Anti-Mouse CD223 (LAG-3) | Real Blue 744 | Becton Dickinson | 756907 | X | |
| Anti-mouse CD366 (Tim-3) Antibody | BV605 | Biolegend | 119721 | X | |
| Rat Anti-Mouse CD19 | BUV615 | Becton Dickinson | 751213 | X | X |
| BD OptiBuild™ Rat Anti-Mouse Siglec-F | BV480 | Becton Dickinson | 746668 | X | X |
| FOXP3 Monoclonal Antibody (FJK-16s) | PE-Cy5 | Invitrogen | 15-5773-80 | X | |
| BD Horizon™ Mouse Anti-Ki-67 | Real Blue 613 | Becton Dickinson | 571127 | X | |
| CD206 (MMR) Monoclonal Antibody (MR6F3) | PE | Invitrogen | 12-2061-82 | | X |
| BD OptiBuild™ Mouse Anti-Mouse NK-1.1 | BV750 | Becton Dickinson | 746876 | | X |
| MHC Class II (I-A/I-E) Monoclonal Antibody (M5/114.15.2) | APC-eFluor780 | Invitrogen | 47-5321-82 | | X |
| Anti-mouse Ly-6C Antibody | Alexa Fluor 488 | Biolegend | 128022 | | X |
| F4/80 Monoclonal Antibody (BM8) | PE-Cyanine5 | Invitrogen | 14-4801-82 | | X |
| BD Horizon™ Rat Anti-Mouse Ly-6G | BUV563 | Becton Dickinson | 612921 | | X |
| Anti-mouse CD86 Antibody | Brilliant Violet (BV) 421 | Biolegend | 105032 | | X |

**Supplementary Table S2. Antibody panel to profile B16OVA wild type tumors.** List of antibodies used for staining of B16OVA tumors for spectral flow cytometry.

| Antibody | Fluorochrome | Company | Catalogue # |
|---|---|---|---|
| BD Horizon™ Rat Anti-Mouse CD31 | BV421 | Becton Dickinson | 562939 |
| Anti-mouse CD45 Antibody | BV650 | Biolegend | 103151 |
| Anti-mouse Podoplanin Antibody | PE-Cy7 | Biolegend | 127412 |
| Anti-mouse CD4 Antibody | Alexa Fluor 647 | Biolegend | 100530 |

**Supplementary Table S3. Antibodies used for bone marrow-derived macrophages (BMDM) activation assay.** List of antibodies used for staining for conventional flow cytometry on BMDM after co-culture with tumor cells.

| BMDM activation | | | |
| --- | --- | --- | --- |
| **Antibody** | **Fluorochrome** | **Company** | **Catalogue #** |
| Anti-Mouse CD45 (30-F11) | cFluor V547 | Cytek | SKU R7-20571 |
| F4/80 Monoclonal Antibody (BM8) | PE-Cyanine5 | Invitrogen | 14-4801-82 |
| Anti-mouse/human CD11b Antibody | Pacific Blue | Biolegend | 101224 |
| CD206 (MMR) Monoclonal Antibody (MR6F3) | PE | Invitrogen | 12-2061-82 |
| Anti-mouse I-A/I-E Antibody (MHC-II) | Alexa Fluor 700 | Biolegend | 107622 |
| Anti-mouse CD86 Antibody | Brilliant Violet (BV) 421 | Biolegend | 105032 |
| CD274 (PD-L1, B7-H1) Monoclonal Antibody (MIH5) | PerCP-eFluor 710 | Invitrogen | 46-5982-80 |

**Supplementary Table S4A.** Relative quantification of GSL-glycan structures assigned based on MS/MS fragmentation and glycobiological pathway constraints. Structures are depicted according to the CFG (Consortium of Functional Glycomics). Blue square: *N*-acetylglucosamine, yellow square: *N*-acetylgalactosamine, blue circle: glucose, yellow circle: galactose, red triangle: fucose, pink diamond: *N*-acetylneuraminic acid, white diamond: *N*-glycolylneuraminic acid. a, b: isomer number; SD: standard variation.

| Glycan number | Glycan name | Proposed structure | Relative abundance %. (SD %) | | Theoretical | Observed | Deviation |
|---|---|---|---|---|---|---|---|
| | | | MOCK | CMAS KO | $[M-H]^-$ | $[M-H]^-$ | $\Delta[M-H]^-$ |
| 1 | Lc2 | 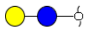 | 0.032 (±0.003) | 0.032 (±0.014) | 343.125 | 343.123 | 0.002 |
| 2 | Gb3 | 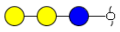 | 2.000 (±0.118) | 85.079 (±2.236) | 505.177 | 505.172 | 0.005 |
| 3 | Gb4 | 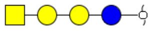 | 0.002 (±0.001) | 0.101 (±0.040) | 708.257 | 708.250 | 0.007 |
| 4 | SSEA3 | 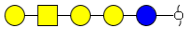 | 0.033 (±0.002) | 0.443 (±0.065) | 870.310 | 870.298 | 0.012 |
| 5 | GM3 | 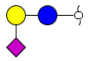 | 0.017 (±0.004) | 0.310 (±0.037) | 634.220 | 634.213 | 0.007 |
| 6 | GM3-Neu5Gc | 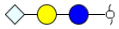 | 0.016 (±0.005) | 0.201 (±0.012) | 650.215 | 650.207 | 0.008 |
| 7 | GM1a | 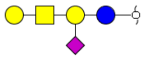 | 0.054 (±0.005) | 0.631 (±0.146) | 999.352 | 999.336 | 0.016 |

| | | | | | | | |
|---|---|---|---|---|---|---|---|
| **8** | **GD1a** | 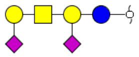 | **96.981 (±0.129)** | **9.409 (±2.198)** | **1290.448** | **1290.427** | **0.021** |
| **9** | **GD3** | 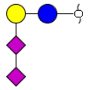 | **0.682 (±0.007)** | **0.949 (±0.162)** | **925.315** | **925.303** | **0.012** |
| **10** | **Gg4** | 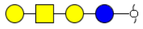 | **0.060 (±0.002)** | **0.925 (±0.163)** | **708.257** | **708.248** | **0.009** |
| **11** | **Gal-Gg4** | 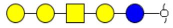 | **0.006 (±0.001)** | **0.171 (±0.030)** | **870.310** | **870.297** | **0.013** |
| **12** | **Lc3** | 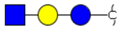 | **0.003 (±0.001)** | **0.021 (±0.004)** | **546.204** | **546.198** | **0.006** |
| **13** | **nLc4** | 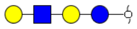 | **0.021 (±0.001)** | **0.198 (±0.028)** | **708.257** | **708.246** | **0.011** |
| **14** | **S(6)nLc4** | 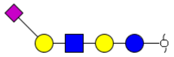 | **0.011 (±0.001)** | **0.087 (±0.014)** | **999.352** | **999.336** | **0.016** |
| **15** | **S(3)nLc4** | 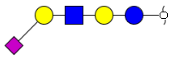 | **0.058 (±0.001)** | **0.923 (±0.107)** | **999.352** | **999.336** | **0.016** |
| **16** | **S(3)nLc4-Neu5Gc** | 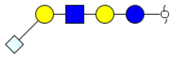 | **0.004 (±0.001)** | **0.052 (±0.005)** | **1015.347** | **1015.326** | **0.021** |
| **17** | **S(3)nLc6** | 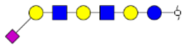 | **0.019 (±0.003)** | **0.187 (±0.019)** | **1364.484** | **1364.484** | **0.000** |

**Supplementary Table S4B.** Relative quantification of O-Glycan structures assigned based on MS/MS fragmentation and glycobiological pathway constraints. Structures are depicted according to the CFG (Consortium of Functional Glycomics). Blue square: *N*-acetylglucosamine, yellow square: *N*-acetylgalactosamine, blue circle: glucose, yellow circle: galactose, red triangle: fucose, pink diamond: *N*-acetylneuraminic acid, white diamond: *N*-glycolylneuraminic acid, S: sulfate. a, b: isomer number; SD: standard variation.

| Glycan number | Glycan composition | Proposed structure | Relative abundance %. (SD %) | | Theoretical | Observed | Deviation |
|---|---|---|---|---|---|---|---|
| | | | MOCK | CMAS KO | $[M-H]^-$ | $\Delta[M-H]^-$ | $\Delta[M-H]^-$ |
| 1 | N1H1 | 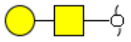 | 0.033 (±0.034) | 5.792 (±0.925) | 384.151 | 384.149 | 0.002 |
| 2 | N1S1 | 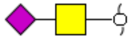 | 1.882 (±0.150) | 0.178 (±0.008) | 513.194 | 513.189 | 0.005 |
| 3 | N1H1S1a | 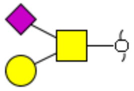 | 3.498 (±0.186) | 0.866 (±0.023) | 675.247 | 675.241 | 0.006 |
| 4 | N1H1S1b | 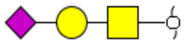 | 14.390 (±0.212) | 5.519 (±0.136) | 675.247 | 675.240 | 0.007 |
| 5 | N1H1S2 | 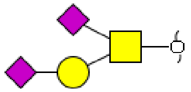 | 58.080 (±1.160) | 1.898 (±0.197) | 966.342 | 966.332 | 0.010 |
| 6 | N2H1 | 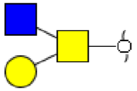 | 0.456 (±0.189) | 9.047 (±0.391) | 587.231 | 587.225 | 0.006 |
| 7 | N2H2a | 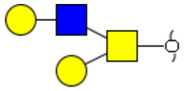 | 0.202 (±0.038) | 45.017 (±2.132) | 749.283 | 749.277 | 0.006 |

| | | | | | | | |
|---|---|---|---|---|---|---|---|
| 8 | N2H2b | 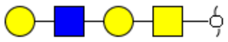 | 0.232 (±0.093) | 1.161 (±0.080) | 749.283 | 749.210 | 0.063 |
| 9 | N2H2F1 | 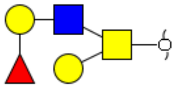 | 0.643 (±0.110) | 3.907 (±0.198) | 895.341 | 895.333 | 0.008 |
| 10 | N2H2F3 | 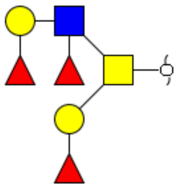 | 1.810 (±0.160) | 5.580 (±1.311) | 1187.457 | 1187.409 | 0.048 |
| 11 | N2H2S2 | 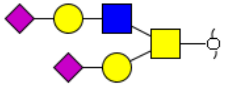 | 11.129 (±0.503) | 2.768 (±0.162) | 1331.474 | 1331.469 | 0.005 |
| 12 | N2H2F1S2 | 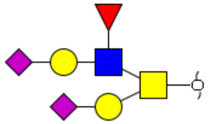 | 0.995 (±0.423) | 3.138 (±0.601) | 1477.532 | 1447.464 | 0.068 |
| 13 | N3H1 | 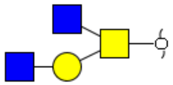 | 3.116 (±0.260) | 10.848 (±0.250) | 790.310 | 790.253 | 0.057 |
| 14 | N3H3 | 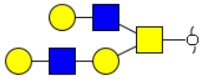 | 0.343 (±0.046) | 2.980 (±0.161) | 1114.416 | 1114.434 | 0.022 |
| 15 | N3H3F1S1Su1 | 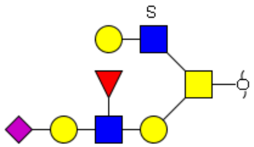 | 2.894 (±0.076) | 1.300 (±0.140) | 1631.526 | 1631.610 | 0.084 |

**Supplementary Table S4C.** Relative quantification of N-Glycan structures assigned based on MS/MS fragmentation and glycobiological pathway constraints. Structures are depicted according to the CFG (Consortium of Functional Glycomics). Blue square: *N*-acetylglucosamine, yellow square: *N*-acetylgalactosamine, blue circle: glucose, yellow circle: galactose, red triangle: fucose, pink diamond: *N*-acetylneuraminic acid, white diamond: *N*-glycolylneuraminic acid, P, phosphate. a, b, c: isomer number; SD: standard variation.

| Glycan number | Glycan composition | Proposed structure | Relative abundance %. (SD %) | | Theoretical | Observed | Deviation |
|---|---|---|---|---|---|---|---|
| | | | MOCK | CMAS KO | $[M-H]^-$ | $[M-H]^-$ | $\Delta[M-H]^-$ |
| 1 | H2N2 | 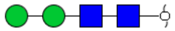 | 0.528 (±0.103) | 0.713 (±0.045) | 749.283 | 749.274 | 0.009 |
| 2 | H2N2F1 | 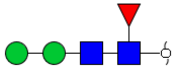 | 0.827 (±0.086) | 1.249 (±0.238) | 895.341 | 895.329 | 0.012 |
| 3 | H3N2 | 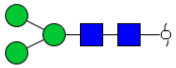 | 0.003 (±0.002) | 0.001 (±0.000) | 911.336 | 911.324 | 0.012 |
| 4 | H3N2P1 | 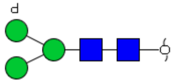 | 0.007 (±0.002) | 0.029 (±0.020) | 991.303 | 991.298 | 0.005 |
| 5 | H3N2F1 | 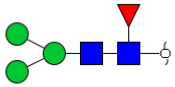 | 0.930 (±0.025) | 3.641 (±0.158) | 1057.394 | 1057.379 | 0.015 |
| 6 | H4N2 | 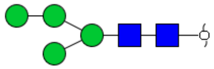 | 0.226 (±0.027) | 0.224 (±0.034) | 1073.389 | 1073.382 | 0.007 |

| | | | | | | | |
|---|---|---|---|---|---|---|---|
| 7 | H4N2F1 | 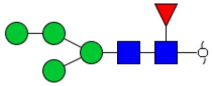 | 0.017(±0.001) | 0.041 (±0.008) | 1219.447 | 1219.441 | 0.006 |
| 8 | H5N2 | 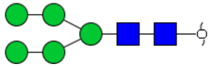 | 1.931 (±0.267) | 1.261 (±0.200) | 1235.442 | 1235.425 | 0.017 |
| 9 | H6N2 | 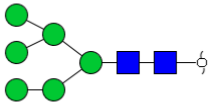 | 8.872 (±0.582) | 8.280 (±0.091) | 1397.495 | 1397.479 | 0.016 |
| 10 | H6N2P1 | 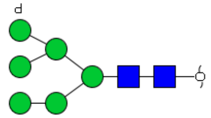 | 1.593 (±0.013) | 1.648 (±0.043) | 1477.461 | 1477.442 | 0.019 |
| 11 | H7N2 | 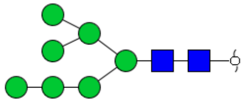 | 6.028 (±0.149) | 6.968 (±0.247) | 1559.547 | 1559.529 | 0.018 |
| 12 | H7N2P1 | 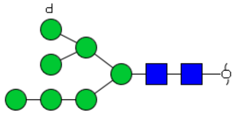 | 1.479 (±0.068) | 1.687 (±0.093) | 1639.514 | 1639.489 | 0.025 |
| 13 | H7N2P2 | 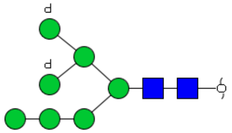 | 0.482 (±0.056) | 1.207 (±0.079) | 1719.480 | 1719.398 | 0.082 |
| 14 | H8N2 | 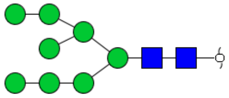 | 11.983 (±0.414) | 13.520 (±1.019) | 1721.600 | 1721.581 | 0.019 |
| 15 | H8N2P1 | 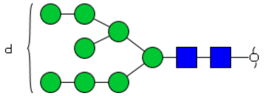 | 0.424 (±0.047) | 0.340 (±0.034) | 1801.567 | 1801.558 | 0.009 |

| | | | | | | | |
|---|---|---|---|---|---|---|---|
| 16 | H9N2 | 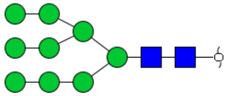 | 14.622 (±0.452) | 16.061 (±1.323) | 1883.653 | 1883.632 | 0.021 |
| 17 | H9N2P1 | 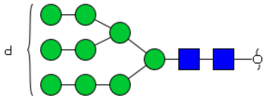 | 0.263 (±0.051) | 0.185 (±0.135) | 1963.619 | 1963.601 | 0.018 |
| 18 | H10N2 | 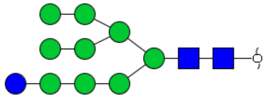 | 1.810 (±0.167) | 2.032 (±0.068) | 2045.706 | 2045.672 | 0.034 |
| 19 | H3N3 | 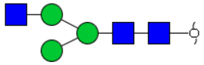 | 0.064 (±0.016) | 0.612 (±0.064) | 1114.416 | 1114.406 | 0.010 |
| 20 | H3N3F1 | 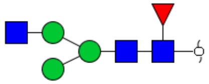 | 0.105 (±0.019) | 1.036 (±0.082) | 1260.473 | 1260.461 | 0.012 |
| 21 | H4N3 | 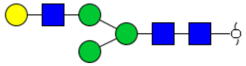 | n.d. | 0.372 (±0.030) | 1276.468 | 1276.459 | 0.009 |
| 22 | H4N3F1 | 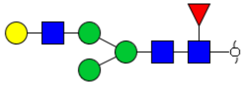 | n.d. | 0.875 (±0.063) | 1422.526 | 1422.514 | 0.012 |
| 23 | H4N3S1 | 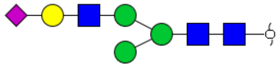 | 0.186 (±0.087) | 0.064 (±0.005) | 1567.564 | 1567.542 | 0.022 |
| 24 | H4N3F1S1 | 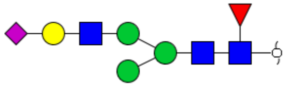 | 0.546 (±0.191) | 0.083 (±0.007) | 1713.622 | 1713.582 | 0.040 |
| 25 | H5N3F1 | 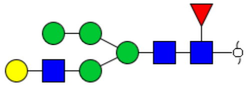 | 0.081 (±0.026) | 0.166 (±0.014) | 1584.579 | 1584.566 | 0.013 |

| | | | | | | | |
|---|---|---|---|---|---|---|---|
| 26 | H5N3S1 | 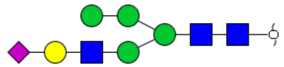 | 0.247 (±0.043) | n.d. | 1729.617 | 1729.609 | 0.008 |
| 27 | H5N3F1S1a | 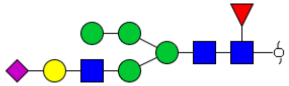 | 0.479 (±0.083) | n.d. | 1875.675 | 1875.640 | 0.035 |
| 28 | H5N3F1S1b | 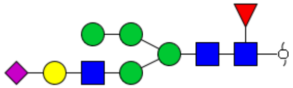 | 0.410 (±0.049) | n.d. | 1875.675 | 1875.642 | 0.033 |
| 29 | H6N3 | 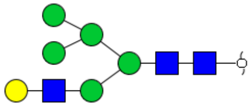 | 0.528 (±0.119) | 0.921 (±0.019) | 1600.574 | 1600.568 | 0.006 |
| 30 | H6N3F1a | 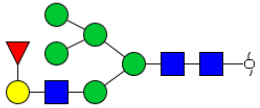 | 0.507 (±0.062) | 0.675 (±0.028) | 1746.632 | 1746.628 | 0.004 |
| 31 | H6N3F1b | 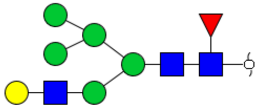 | 0.165 (±0.073) | 0.381 (±0.039) | 1746.632 | 1746.626 | 0.006 |
| 32 | H6N3S1a | 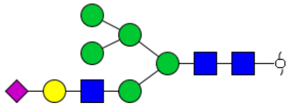 | 1.021 (±0.159) | n.d. | 1891.669 | 1891.637 | 0.032 |
| 33 | H6N3S1b | 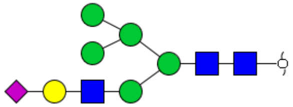 | 0.985 (±0.149) | n.d. | 1891.669 | 1891.635 | 0.034 |
| 34 | H6N3F1S1a | 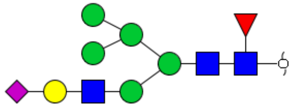 | 0.398 (±0.072) | n.d. | 2037.727 | 2037.693 | 0.034 |

| | | | | | | | |
|---|---|---|---|---|---|---|---|
| 35 | H6N3F1S1b | 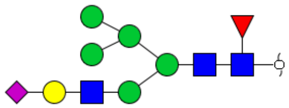 | 0.208 (±0.038) | n.d. | 2037.727 | 2037.693 | 0.034 |
| 36 | H7N3F1a | 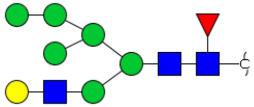 | 0.659 (±0.070) | 0.851 (±0.031) | 1908.685 | 1908.678 | 0.007 |
| 37 | H7N3F1b | 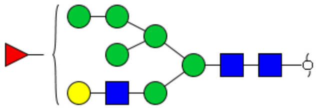 | 0.129 (±0.092) | 0.301 (±0.099) | 1908.685 | 1908.676 | 0.009 |
| 38 | H3N4 | 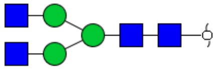 | 0.085 (±0.007) | 0.179 (±0.012) | 1317.495 | 1317.486 | 0.009 |
| 39 | H3N4F1 | 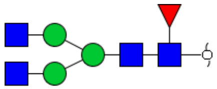 | 0.335 (±0.039) | 0.590 (±0.042) | 1463.553 | 1463.533 | 0.020 |
| 40 | H4N4F1 | 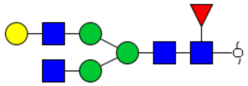 | 0.306 (±0.076) | 2.741 (±0.083) | 1625.606 | 1625.577 | 0.029 |
| 41 | H4N4F1S1 | 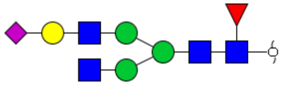 | 0.468 (±0.063) | 0.086 (±0.035) | 1916.701 | 1916.667 | 0.034 |
| 42 | H5N4 | 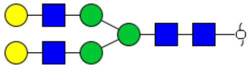 | 0.161 (±0.133) | 2.673 (±0.255) | 1641.601 | 1641.568 | 0.033 |
| 43 | H5N4F1 | 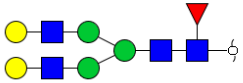 | 0.501 (±0.336) | 9.241 (±0.168) | 1787.659 | 1787.626 | 0.033 |

| | | | | | | | |
|---|---|---|---|---|---|---|---|
| 44 | H5N4S1a | 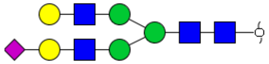 | 1.102 (±0.269) | 0.132 (±0.008) | 1932.696 | 1932.662 | 0.034 |
| 45 | H5N4S1b | 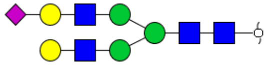 | 0.935 (±0.286) | 0.115 (±0.014) | 1932.696 | 1932.661 | 0.035 |
| 46 | H5N4F1S1a | 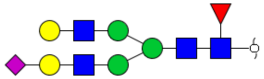 | 1.078 (±1.295) | 0.016 (±0.007) | 2078.754 | 2078.719 | 0.034 |
| 47 | H5N4F1S1b | 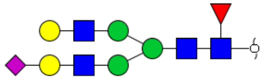 | 1.413 (±0.784) | 0.102 (±0.017) | 2078.754 | 2078.720 | 0.034 |
| 48 | H5N4S2a | 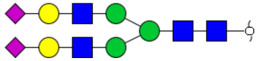 | 0.762 (±0.114) | n.d. | 2223.791 | 2223.753 | 0.038 |
| 49 | H5N4S2b | 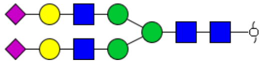 | 1.748 (±0.418) | n.d. | 2223.791 | 2223.754 | 0.037 |
| 50 | H5N4S2c | 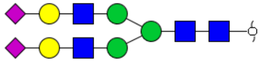 | 0.569 (±0.145) | n.d. | 2223.791 | 2223.754 | 0.037 |
| 51 | H5N4F1S2a | 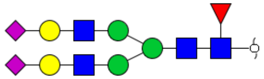 | 2.133 (±0.315) | n.d. | 2369.849 | 2369.819 | 0.030 |
| 52 | H5N4F1S2b | 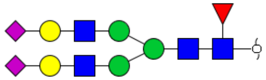 | 4.315 (±0.1.020) | n.d. | 2369.849 | 2369.811 | 0.038 |
| 53 | H5N4F1S2c | 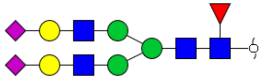 | 0.974 (±0.340) | n.d. | 2369.849 | 2369.810 | 0.039 |
| 54 | H5N5 | 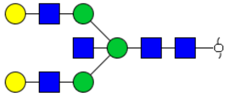 | n.d. | 0.554 (±0.043) | 1844.680 | 1844.643 | 0.037 |

| | | | | | | | |
|---|---|---|---|---|---|---|---|
| 55 | H5N5F1a | 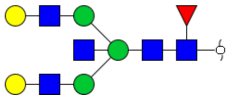 | n.d. | 1.248 (±0.085) | 1990.738 | 1990.710 | 0.028 |
| 56 | H5N5F1b | 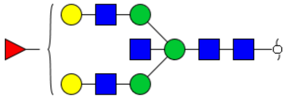 | n.d. | 0.401 (±0.034) | 1990.738 | 1990.697 | 0.041 |
| 57 | H5N5S1a | 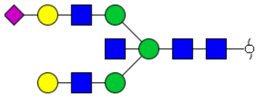 | 0.156 (±0.008) | n.d. | 2135.775 | 2135.737 | 0.038 |
| 58 | H5N5S1b | 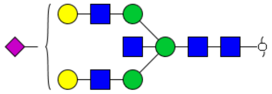 | 0.095 (±0.029) | n.d. | 2135.775 | 2135.738 | 0.037 |
| 59 | H5N5S2 | 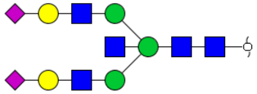 | 0.265 (±0.085) | n.d. | 2426.871 | 2426.829 | 0.042 |
| 60 | H6N5a | 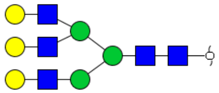 | n.d. | 2.205 (±0.123) | 2006.733 | 2006.696 | 0.037 |
| 61 | H6N5b | 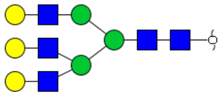 | n.d. | 1.532 (±0.146) | 2006.733 | 2006.696 | 0.037 |
| 62 | H6N5F1a | 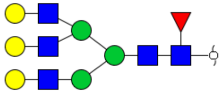 | 0.074 (±0.043) | 5.991 (±0.799) | 2152.791 | 2152.752 | 0.039 |
| 63 | H6N5F1b | 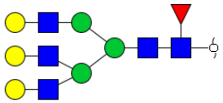 | 0.074 (±0.042) | 3.488 (±0.048) | 2152.791 | 2152.749 | 0.042 |

| | | | | | | | |
|---|---|---|---|---|---|---|---|
| 64 | H6N5F1c | 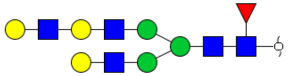 | n.d. | 0.516 (±0.050) | 2152.791 | 2152.750 | 0.041 |
| 65 | H6N5F1S1a | 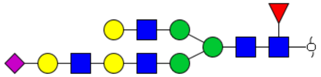 | 0.134 (±0.071) | 0.081 (±0.013) | 2443.886 | 2443.841 | 0.045 |
| 66 | H6N5F1S1b | 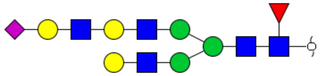 | 0.228 (±0.108) | 0.078 (±0.007) | 2443.886 | 2443.842 | 0.044 |
| 67 | H6N5F1S1c | 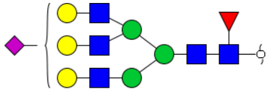 | 0.176 (±0.107) | n.d. | 2443.886 | 2443.842 | 0.044 |
| 68 | H5N5F1S2 | 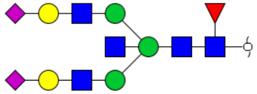 | 1.440 (±0.688) | n.d. | 2572.929 | 2572.890 | 0.039 |
| 69 | H6N5F1S2 | 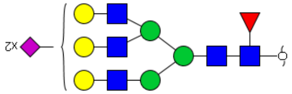 | 5.802 (±2.619) | n.d. | 2734.982 | 2734.749 | 0.033 |
| 70 | H6N5S3 | 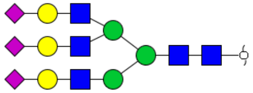 | 1.124 (±0.211) | n.d. | 2880.019 | 2879.970 | 0.049 |
| 71 | H6N5F1S3a | 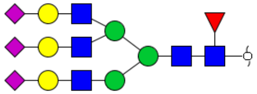 | 0.602 (±0.243) | n.d. | 3026.077 | 3026.025 | 0.052 |
| 72 | H6N5F1S3b | 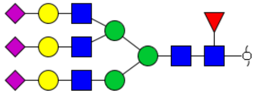 | 2.688 (±0.936) | n.d. | 3026.077 | 3026.027 | 0.050 |

| | | | | | | | |
|---|---|---|---|---|---|---|---|
| **73** | **H7N6F1** | 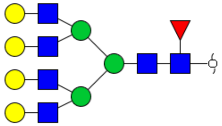 | **0.084 (±0.020)** | **2.603 (±0.282)** | **2517.923** | **2517.892** | **0.031** |
| **74** | **H7N6F1S2** | 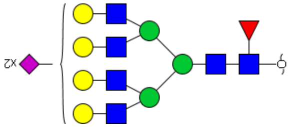 | **1.554 (±0.511)** | **n.d.** | **3100.114** | **3100.056** | **0.058** |
| **75** | **H7N6F1S3** | 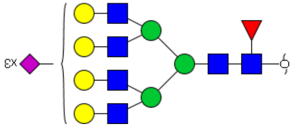 | **3.095 (±0.660)** | **n.d.** | **3391.209** | **3391.137** | **0.072** |
| **76** | **H7N6S4** | 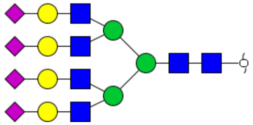 | **0.916 (±0.469)** | **n.d.** | **3536.247** | **3536.194** | **0.053** |
| **77** | **H7N6F1S4** | 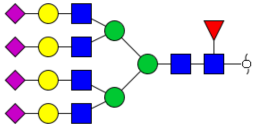 | **5.865 (±3.977)** | **n.d.** | **3682.305** | **3682.251** | **0.054** |
